# Sexual lineage specific DNA methylation regulates Arabidopsis meiosis

**DOI:** 10.1101/201798

**Authors:** James Walker, Hongbo Gao, Jingyi Zhang, Billy Aldridge, Martin Vickers, James D. Higgins, Xiaoqi Feng

**Affiliations:** Department of Cell and Developmental Biology, John Innes Centre, Norwich NR4 7UH, UK.; Department of Genetics, University of Leicester, University Road, Leicester LE1 7RH, UK.

**Author notes:** These authors contributed equally to this work (James Walker, Hongbo Gao). For correspondence: Xiaoqi Feng.

## Abstract

DNA methylation controls eukaryotic gene expression and is extensively reprogrammed to regulate animal development. However, whether developmental methylation reprogramming during the sporophytic life cycle of flowering plants regulates genes is presently unknown. Here we report a distinctive, gene-targeted RNA-directed DNA methylation (RdDM) activity in the *Arabidopsis thaliana* male sexual lineage that regulates gene expression in meiocytes. Loss of sexual lineage-specific RdDM causes mis-splicing of the *MPS1/PRD2* gene, thereby disrupting meiosis. Our results establish a regulatory paradigm in which *de novo* methylation creates a cell-lineage-specific epigenetic signature that controls gene expression and contributes to cellular function in flowering plants.

## MAIN TEXT

Cytosine methylation is an ancient DNA modification catalyzed by methyltransferases that are conserved across eukaryotes, including plants and animals^1^. Cytosine methylation in the CG dinucleotide context is maintained by DNA Methyltransferase 1 (Dnmt1, called MET1 in plants), which recognizes hemimethylated CG dinucleotides and adds a methyl group to the unmethylated cytosine during DNA replication^2,3^. Plant methylation also commonly occurs in the CHG and CHH (H=A, C, or T) contexts, maintained by the CHROMOMETHYLASE3 (CMT3) and CMT2 enzymes, respectively^4,5^. Establishment of *de novo* methylation is catalyzed by Dnmt3 and its plant DRM homologues^2,3^. DRM enzymes (DRM1 and 2 in *Arabidopsis thaliana*) are part of the RNA-directed DNA methylation (RdDM) pathway, which typically targets transposons and methylates cytosines regardless of sequence context^6^. In RdDM, 24-nucleotide (nt) small RNAs (sRNAs), which are produced from transcripts created by plant-specific RNA polymerase IV and RNA-dependent RNA polymerase 2 (RDR2), guide DRM methyltransferases to DNA via association with a homologous transcript generated by RNA polymerase V and the DRD1 chromatin remodeling protein^5,6^.

DNA methylation patterns are faithfully replicated during cell division, which allows methylation to carry epigenetic information within cellular lineages^2,3^. In the complex genomes of flowering plants and vertebrates, methylation heritably silences transposons, thereby maintaining genome integrity and transcriptional homeostasis^2,3^. Consistent with this function, DNA methylation of regulatory sequences, especially those near transcriptional start sites, is strongly associated with gene silencing^5,7^.

In addition to its homeostatic function, DNA methylation can be reprogrammed during development to regulate gene expression. In mammals, this phenomenon has been observed in a number of tissues and cellular lineages, and appears to be a common regulatory mechanism^8–14^. In plants, gene expression in the transient extra-embryonic endosperm tissue is controlled by active DNA demethylation, which occurs in the central cell (a companion cell of the egg) that gives rise to the endosperm^15,16^. A similar active demethylation process also occurs in the vegetative cell, a terminally differentiated companion cell of the sperm^15,17,18^. Beyond the endosperm and gamete companion cells, although intriguing examples of altered methylation levels and patterns in different cell types^17–20^ and during responses to biotic and abiotic stimuli ^21–25^ have been described, whether gene expression is controlled by developmental reprogramming of DNA methylation in plants is unknown.

To investigate this question, we analyzed DNA methylation in the male sexual lineage of *Arabidopsis thaliana*. This allowed us to uncover a sexual-lineage specific DNA methylation signature deposited by the RdDM pathway. We further demonstrated that this *de novo* methylation regulates gene expression and splicing, and is required for normal meiosis, providing a compelling link between DNA methylation reprogramming, gene expression and developmental outcome. The RdDM pathway is widely present in plant tissues, and therefore has the potential to regulate the development of many cell types and tissues.

### Male meiocytes feature a typical germline methylome with high CG and low CHH methylation

In *Arabidopsis* and other flowering plants, the male sexual lineage initiates as diploid meiocytes, which give rise to haploid microspores via meiosis^26^ (Fig. 1a). The microspores subsequently divide mitotically to produce the vegetative and generative cells (Fig. 1a). The generative cell enters one more round of mitosis to generate two sperm cells, which are engulfed within the vegetative cell in the mature pollen grain (Fig. 1a). To comprehensively understand DNA methylation reprogramming within the entire lineage, we generated a genome-wide methylation profile for male *Arabidopsis thaliana* meiocytes (Supplementary Table 1), which we compared to those of the microspore, sperm and vegetative cell^17,18^.

**Figure 1.**
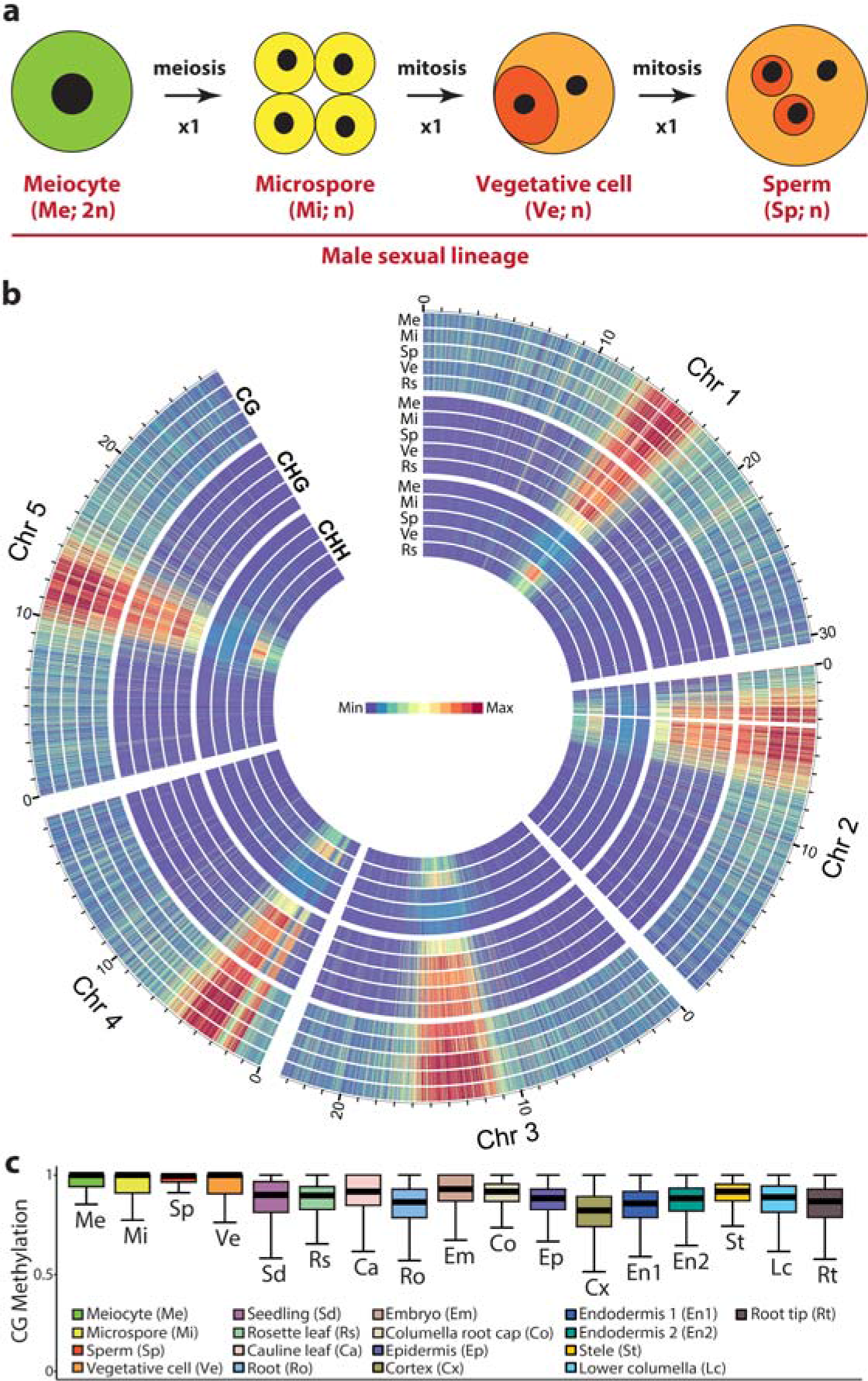
Male meiocytes exhibit high CG/CHG and low CHH methylation. **a**, Model of male sexual lineage development in *Arabidopsis thaliana* n, the number of chromosomes in the haploid genome. **b**, Heat maps showing CG, CHG and CHH methylation of the male sexual lineage comprising the meiocyte (Me), microspore (Mi), sperm (Sp) and vegetative cell (Ve), in comparison to rosette leaf (Rs). Methylation is calculated and presented in 10 kb windows, with the maximum set at the highest value among the five tissues for each context. The region enriched with mitochondria sequences on Chromosome 2 (3.23 to 3.51 Mb) is removed. **c**, Box plots demonstrating CG methylation for individual CG sites located within annotated transposons, and with methylation greater than 50% and at least 10 informative sequenced cytosines. Each box encloses the middle 50% of the distribution, with the horizontal line making the median, and vertical lines marking the minimum and maximum values that fall within 1.5 times the height of the box.

Contrary to the speculation of DNA demethylation in male meiocytes^27^, we found meiocyte methylation resembles that of microspores and sperm, with high levels of CG and CHG methylation in transposons (Fig. 1b,c and Supplementary Fig. 1). This is consistent with robust transposon silencing in the germline, an essential function for ensuring genetic integrity across generations^16,28^. In the CHH context, the microspore and sperm cells of the germline have low levels of methylation compared to somatic tissues and, especially, vegetative cells^17,18,28^ (Fig. 1b and Supplementary Fig. 1). However, the male meiocyte has even lower CHH methylation than microspore and sperm cells (Fig. 1b and Supplementary Fig. 1). Low levels of CHH methylation in microspore and sperm were proposed to result from lack of methylation maintenance during meiotic divisions^17^. Our results demonstrate this is not the case, and that CHH methylation instead undergoes an overall increase during the development of the male sexual lineage.

### Hypermethylated loci are observed in the male sexual lineage

Comparison of DNA methylation patterns between male sex cells and somatic tissues (seedlings, rosette leaves, cauline leaves, roots) revealed regions that are strongly hypermethylated in sex cells (Fig. 2a and Supplementary Fig. 2). Furthermore, loci hypermethylated in one male sex cell type tend to be hypermethylated in other sex cells (Fig. 2a,b and Supplementary Fig. 2). This hypermethylation is most prominent in the CHH context, but encompasses other contexts as well (Fig. 2a,b and Supplementary Fig. 2), so that the same locus is often hypermethylated at CG, CHG and CHH sites (Fig. 2a and Supplementary Fig. 2). Among the 1301 loci we identified that are consistently differentially methylated between sex cells and somatic tissues, the overwhelming majority (1265; 97%) are hypermethylated in sex cells (Supplementary Data 1). These sexual-lineage hypermethylated loci (SLHs) are typically small (529 nucleotides on average; Supplementary Data 1), altogether encompassing 0.6% of the nuclear genome.

**Figure 2.**
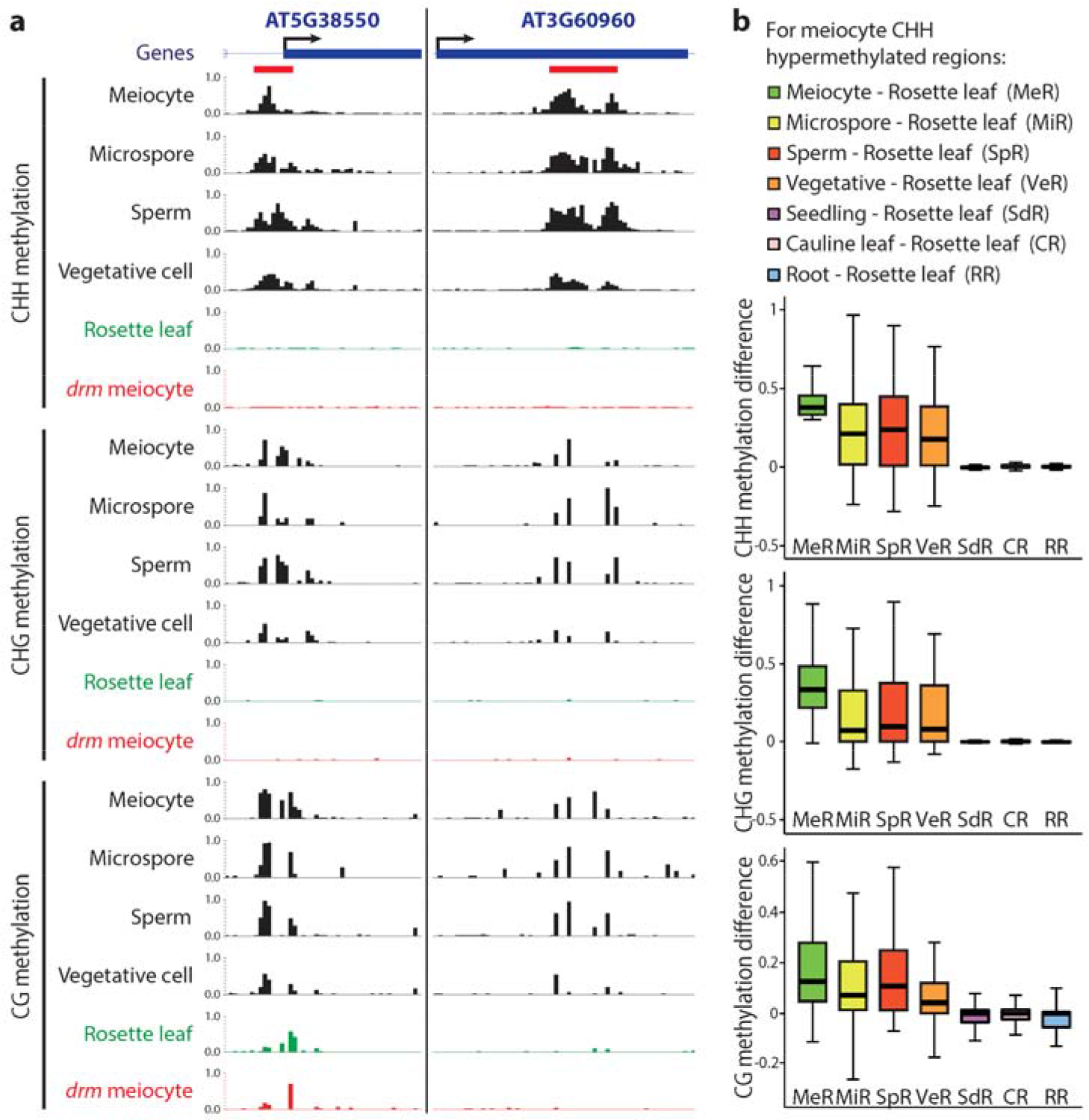
Sexual-lineage hypermethylated loci (SLHs) in *Arabidopsis*. **a**, Snapshots of cytosine methylation in wild-type male sex cells (black), rosette leaf (green) and *drm1drm2* (*drm*) mutant meiocyte (red) at two SLH examples. SLHs (refer to Supplementary Data 1 for a full list) are underlined in red. Methylation patterns in other somatic tissues and *drm* sex cells are shown in Supplementary Fig. 2. **b**, Box plots showing absolute methylation difference between specific cells/tissues and rosette leaf for 50 bp windows that are CHH hypermethylated in meiocytes in comparison to rosette leaves.

### Sexual-lineage hypermethylated loci (SLHs) are caused by RdDM

The SLHs resemble targets of the small RNA-directed DNA methylation (RdDM) pathway, which establishes and maintains methylation in all sequence contexts, but is particularly important for CHH methylation of relatively small loci^3,5,6^. To test whether RdDM is responsible for SLHs, we analyzed the methylomes of meiocytes, sperm and vegetative cells with mutations in both *DRM1* and *DRM2*, as well as the methylomes of sperm and vegetative cells with a mutation in *RDR2* (Supplementary Table 1). In all examined mutant sex cells, SLHs are extensively hypomethylated in all sequence contexts (Figs 2a and 3a; Supplementary Fig. 2), demonstrating that SLHs are a product of RdDM. As expected of RdDM targets in the sexual lineage, SLHs are associated with the 24-nt sRNAs that guide RdDM at levels similar to those of other RdDM target loci in pollen, but not in shoots (Fig. 3b). The vast majority of SLHs (99.4%; 1257 loci) have significantly less methylation in *drm1drm2* mutant sex cells in comparison to those of wild type (none in the reverse comparison; both *P* < 0.001; Fisher’s exact test); this slightly reduced group of SLHs is used for subsequent analyses (Supplementary Data 1). Collectively, our results demonstrate that RdDM is the underlying mechanism producing SLHs.

In *Arabidopsis*, RdDM is counteracted by active DNA demethylation^3^, suggesting that somatic demethylation may contribute to the distinct patterns of RdDM in male sex cells and somatic tissues. To test this possibility, we examined available methylation data for rosette leaves with mutations in the three demethylases expressed in somatic tissues: *ROS1*, *DML2* and *DML3*^29^. CG methylation at SLHs in these *rdd* mutants is indeed much higher than in wild-type control leaves, but not as high as in wild-type sex cells (Fig. 3c). CHG methylation in *rdd* leaves is also higher than in wild-type, but substantially lower than in sex cells (Fig. 3c), and CHH methylation is only slightly higher in *rdd* leaves compared to wild-type and much lower than in sex cells (Fig. 3c). Given the known ability of RdDM-established CG methylation, but not CHH methylation, to be maintained in the absence of RdDM^6^, our data suggest that active demethylation removes some of the CG (and CHG) methylation that is induced by RdDM in the sexual lineage and maintained in somatic tissues.

**Figure 3.**
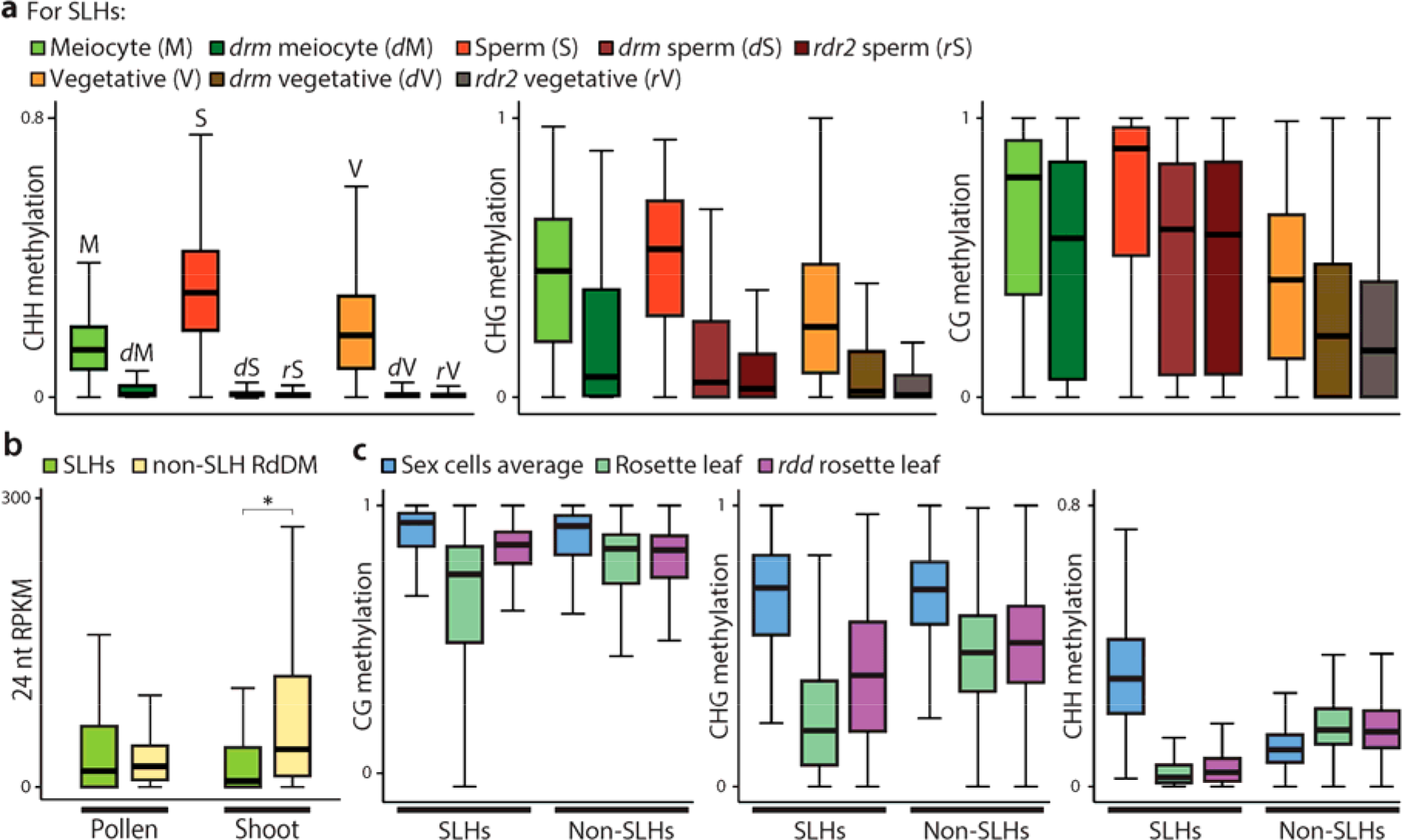
Sexual-lineage hypermethylated loci (SLHs) are produced by RdDM. **a**, Box plots showing the absolute methylation at SLHs in the *drm1drm2* (*drm*) and *rdr2* mutants in comparison to wild type. **b**, Box plots demonstrating the abundance of 24 nucleotide (nt) small RNA in pollen or shoot at SLHs and non-SLH RdDM target loci. \**P* < 0.001; Kolmogorov-Smirnov test. **c**, Box plots showing the absolute methylation at SLH and non-SLH 50 bp windows in *ros1dml2dml3* (*rdd*) mutant rosette leaf, wild-type rosette leaf, and wild-type male sex cells (meiocyte, microspore and sperm).

### Many SLHs are novel RdDM targets specific to the sexual lineage

As a manifestation of RdDM activity in the sexual lineage, SLHs can simply be a product of increased RdDM activity at canonical targets, an expansion of RdDM into novel targets, or both. As CHH/G methylation levels are indicators of RdDM activity, we separated SLHs into two groups based on the level of CHH/G methylation in somatic tissues: canonical SLHs (724 loci), which have CHH/G methylation in the soma, and sexual-lineage specific methylated loci (SLMs; 533 loci), which lack CHH/G methylation in somatic tissues (Supplementary Data 1).

To further evaluate the cell/tissue specificity of SLMs, we examined the root cap columella cell, which has high levels of RdDM-associated CHH methylation^19^. Whereas canonical SLHs show methylation in all sequence contexts in columella cells (Fig. 4a), SLMs have little CHH/G methylation in columella (Supplementary Fig. 3). Furthermore, whereas 76% (551/724) of the canonical SLHs overlap with published columella differentially methylated regions (DMRs)^19^, most SLMs (88%, 469 sites; Supplementary Data 1) do not overlap with columella DMRs and show no CHH/G methylation in columella (Fig. 4b). Examination of DNA methylation in the embryo – another tissue with reported CHH hypermethylation^20^ – revealed that although canonical SLHs are methylated (Fig. 4a), the 469 SLMs that do not overlap with columella DMRs show no CHH/G methylation in embryo (Fig. 4b). We use this group of highly specific SLMs for all subsequent analyses (Supplementary Data 1).

**Figure 4.**
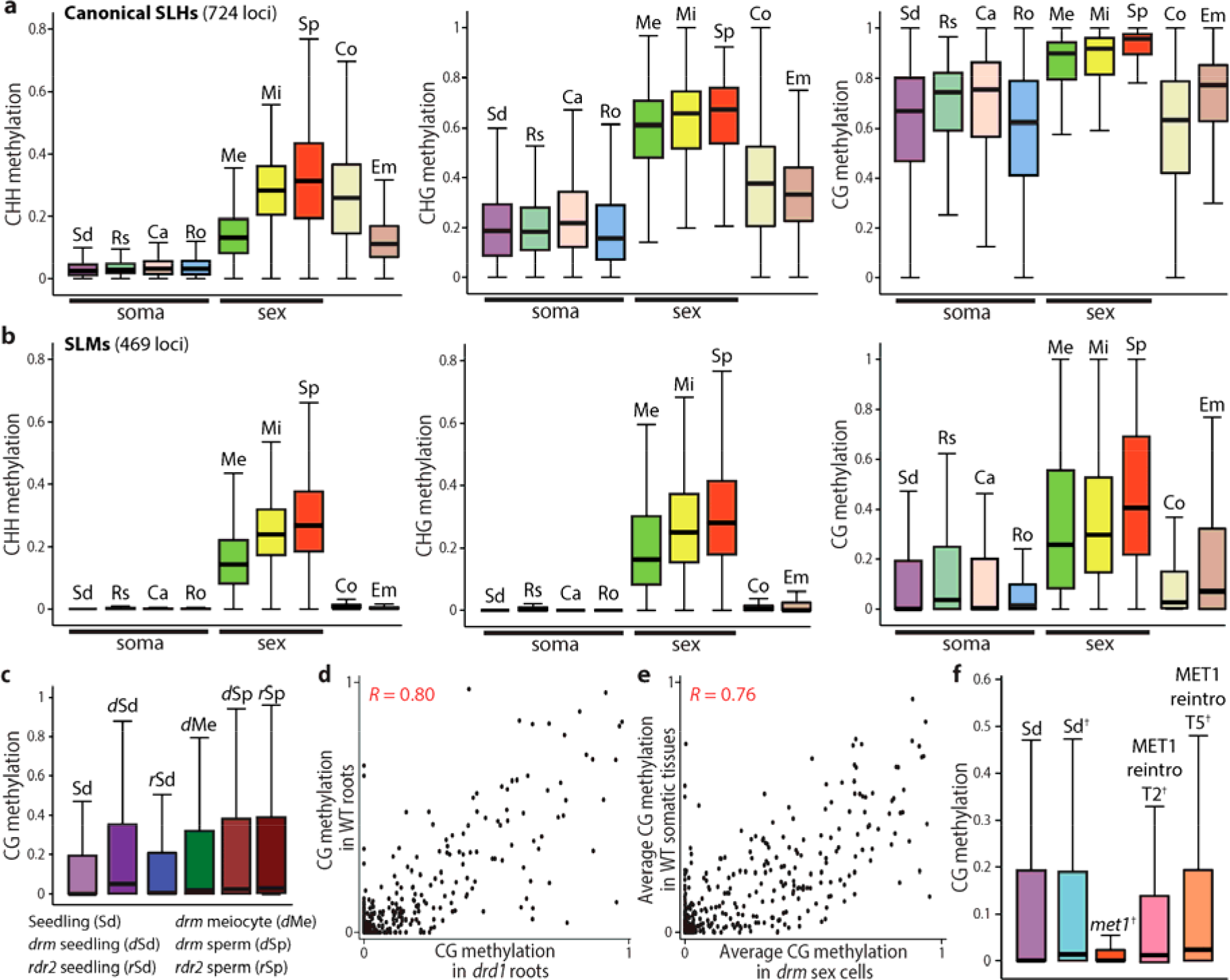
SLMs are novel RdDM targets specific to the sexual lineage. **a,b,** Box plots showing the absolute methylation at canonical SLHs (a) and SLMs (b) in somatic tissues (Sd, seedling; Rs, rosette leaf; Ca, cauline leaf; Ro, root), sex cells (Me, meiocyte; Mi, microspore; Sp, sperm), columella root cap (Co) and embryo (Em). **c**, Box plots illustrating CG methylation at SLMs in wild-type (WT) seedling (refer to b for other somatic tissues), and seedlings and sex cells from RdDM mutants: *drm1drm2* (*drm*) and *rdr2*. **d**, Scatter plot showing the linear correlation between CG methylation in WT (*y* axis) and *drdl* mutant (*x* axis) roots at SLMs (Pearson’s *R* = 0.80). **e**, Scatter plot showing the linear correlation between average CG methylation in WT somatic tissues (cauline leaf, rosette leaf, root and seedling; *y* axis) and that in *drm* mutant sex cells (meiocytes, sperm and vegetative cell; *x* axis) at SLMs (Pearson’s *R* = 0.76). **f**, Box plots demonstrating the absolute CG methylation at SLMs in WT seedling (same data as used in b and c), and published data^†30^ including WT control seedlings (Sd^†^), *metl* mutant, and *MET1* reintroduction lines (T-MET1a T2 and T-MET1b T5).

Although SLMs lack CHH/G methylation in somatic tissues, some CG methylation is present (Fig. 4b and Supplementary Fig. 2d,e). This remnant CG methylation can either be induced by sexual-lineage specific RdDM and maintained in somatic tissues by MET1, or result directly from somatic RdDM activity. To distinguish between these hypotheses, we analyzed SLM CG methylation in RdDM mutant somatic tissues, which showed similar overall levels to wild-type somatic tissues (Fig. 4c). Furthermore, SLM CG methylation in RdDM (*drd1*, *drm2* and *rdr2*) mutant somatic tissues correlates with that in wild-type (Pearson’s *R* = 0.80, 0.58 and 0.70, respectively; Fig. 4d and Supplementary Fig. 4), demonstrating that RdDM is not required to maintain somatic CG methylation at SLMs.

The hypothesis that CG methylation at SLMs is initiated by RdDM in sex cells and is maintained at lower levels by MET1 in the absence of RdDM in somatic cells makes several predictions. First, MET1 should be able to maintain CG methylation in sex cells without RdDM at levels similar to wild-type somatic tissues. Indeed, SLM CG methylation in *drm1drm2* mutant sex cells is similar to wild-type somatic tissues (Fig. 4c and Supplementary Fig. 2d,e) and strongly correlates with that in wild-type somatic tissues (Pearson’s *R* = 0.76; Fig. 4e). Second, somatic CG methylation at SLMs should be MET1-dependent, which it is (Fig. 4f). Finally, CG methylation at SLMs should be reestablished after it is erased, because CG methylation is known to be reestablished at loci that are targeted by RdDM in a manner that is not dependent on preexisting CG methylation (*i.e*. at loci where RdDM still works in *met1* mutants)^30^. Indeed, somatic CG methylation at SLMs is restored to wild-type levels through introduction of functional MET1 into *met1* mutant plants (Fig. 4f). Taken together, our analyses demonstrate that SLMs are products of sexual-lineage specific RdDM activity, which establishes methylation in all sequence contexts. In somatic tissues, residual CG methylation at SLMs is maintained by MET1 in the absence of RdDM.

### RdDM-induced sexual-lineage specific methylation regulates gene expression in meiocytes

Because SLMs are not targeted by RdDM outside the sexual lineage, we analyzed whether they resemble conventional RdDM loci by comparing the proximity of SLMs, canonical SLHs, and other RdDM targets to genes and transposons. Canonical SLHs correspond mostly to transposons, but overlap genes more frequently than other RdDM targets (Fig. 5a). Furthermore, canonical SLHs are more likely to overlap annotated transposons than randomly selected sets of loci that are comparably located in relation to genes throughout the genome, but are less likely to overlap transposons than other RdDM target loci (Supplementary Table 2). Surprisingly, the majority of SLMs overlap genes (Fig. 5a), and are even slightly less likely to overlap annotated transposons than random control loci (Supplementary Table 2). These results indicate that canonical SLHs are an extension of conventional, transposon-targeted RdDM, which is consistent with their methylation in some somatic cell types, whereas SLMs are an aggressive expansion of RdDM into genes.

**Figure 5.**
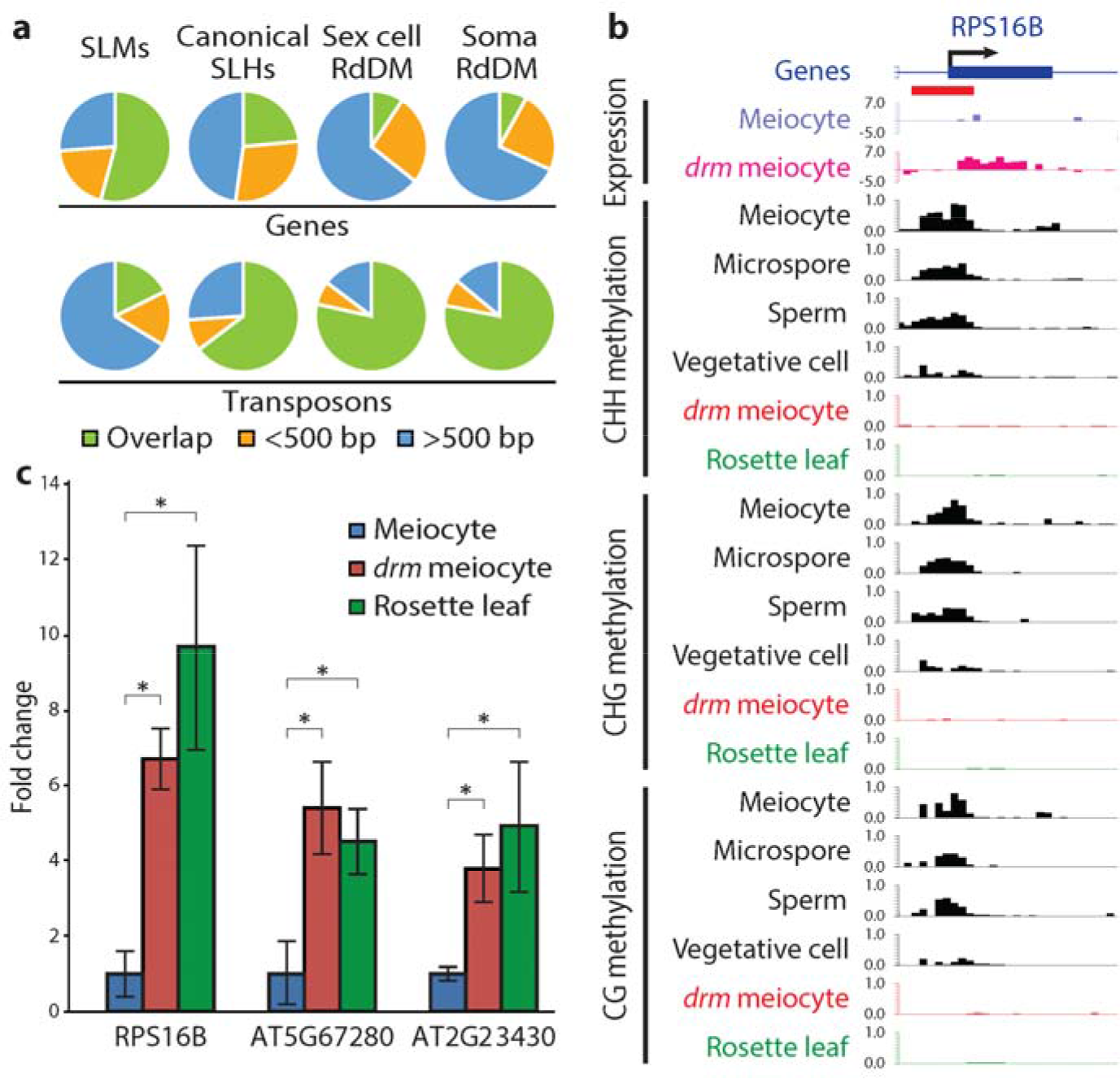
SLMs target genes and regulate gene expression in meiocytes. **a**, Pie charts illustrating percentages of SLMs, canonical SLHs and other RdDM target loci overlapping (green), within 500 bp (yellow), and more than 500 bp from (blue) genes or transposons (numbers shown in Supplementary Table 2). **b**, Snapshots of transcription (in log_2_RPKM) and DNA methylation (similar to Fig. 2a), at the *RPS16B* gene. *drm*, *drm1drm2*; SLM is underlined in red. **c**, Quantitative RT-PCR showing the expression of three SLM-regulated genes. \**P* < 0.02 (*t*-test).

The common occurrence of SLMs in genes and the well-demonstrated role of DNA methylation in suppressing gene expression^3^ suggested that SLMs may repress gene expression in the sexual lineage. To test this hypothesis, we analyzed mRNA levels in *drm1drm2* mutant meiocytes and wild-type controls by RNA-seq. Expression of meiosis-associated genes is substantially enriched in our data compared with published meiocyte RNA-seq results (Supplementary Data 2)^31,32^, suggesting high meiocyte purity. Among the 47 genes with a greater than four-fold change in expression between wild-type and *drm1drm2* meiocytes, all of which are activated in *drm1drm2* meiocytes, seven overlap an SLM and one has an SLM within 20 bp (Fig. 5b,c, Supplementary Fig. 5 and Supplementary Data 3), a much higher fraction (17%) than expected by random chance (Fisher’s exact test, *P* = 1.40х10^−8^), as only 0.9% of nuclear genes are within 20 bp of an SLM. Furthermore, all four of the SLM-associated genes that are overexpressed in *drm1drm2* meiocytes and significantly differentially expressed between meiocytes and leaves are suppressed in meiocytes compared to leaves (Fig. 5c and Supplementary Data 3). Expression levels of these genes in leaves are not elevated by RdDM mutation (Supplementary Data 3). These data indicate that RdDM-mediated SLMs specifically regulate gene expression in meiocytes.

### Pre-tRNA genes encoding specific anti-codons are hypermethylated in the male sex lineage

Among the genes containing SLMs, an unexpected group is comprised of pre-tRNA genes. 24 pre-tRNA loci overlap SLMs, with preference for specific anti-codons: for example, 75% and 21% of the phenylalanine and methionine pre-tRNA genes are covered by SLMs (Fig. 6a, Supplementary Fig. 6 and Supplementary Table 3a); numbers that are substantially higher than expected by random chance (both *P* < 2.63х10^−6^, Fisher’s exact test). As our criteria for calling SLMs are very stringent, we performed a genome-wide analysis to specifically detect sexual lineage hypermethylation of pre-tRNA genes. We found an additional set of 60 pre-tRNAs with significantly more CHH and CHG methylation in at least two of the sex cells in comparison to somatic tissues, and in wild-type sex cells in comparison to *drm1drm2* sex cells (both *P* < 0.001, Fisher’s exact test; Fig. 6b, Supplementary Fig. 7 and Supplementary Table 3b). Together, the 84 hypermethylated loci include 100%, 75%, 73% and 42% of the phenylalanine, valine, cysteine and methionine pre-tRNA genes, respectively (Supplementary Table 3b). Consistently, 24-nt sRNAs are enriched at these pre-tRNA genes in pollen, but not in shoots (Fig. 6c). The preferential hypermethylation of certain pre-tRNA genes, together with the recent discovery of small tRNA fragments in *Arabidopsis* pollen^33^, suggest that tRNA biology may have interesting aspects that are particular to sex cells.

**Figure 6.**
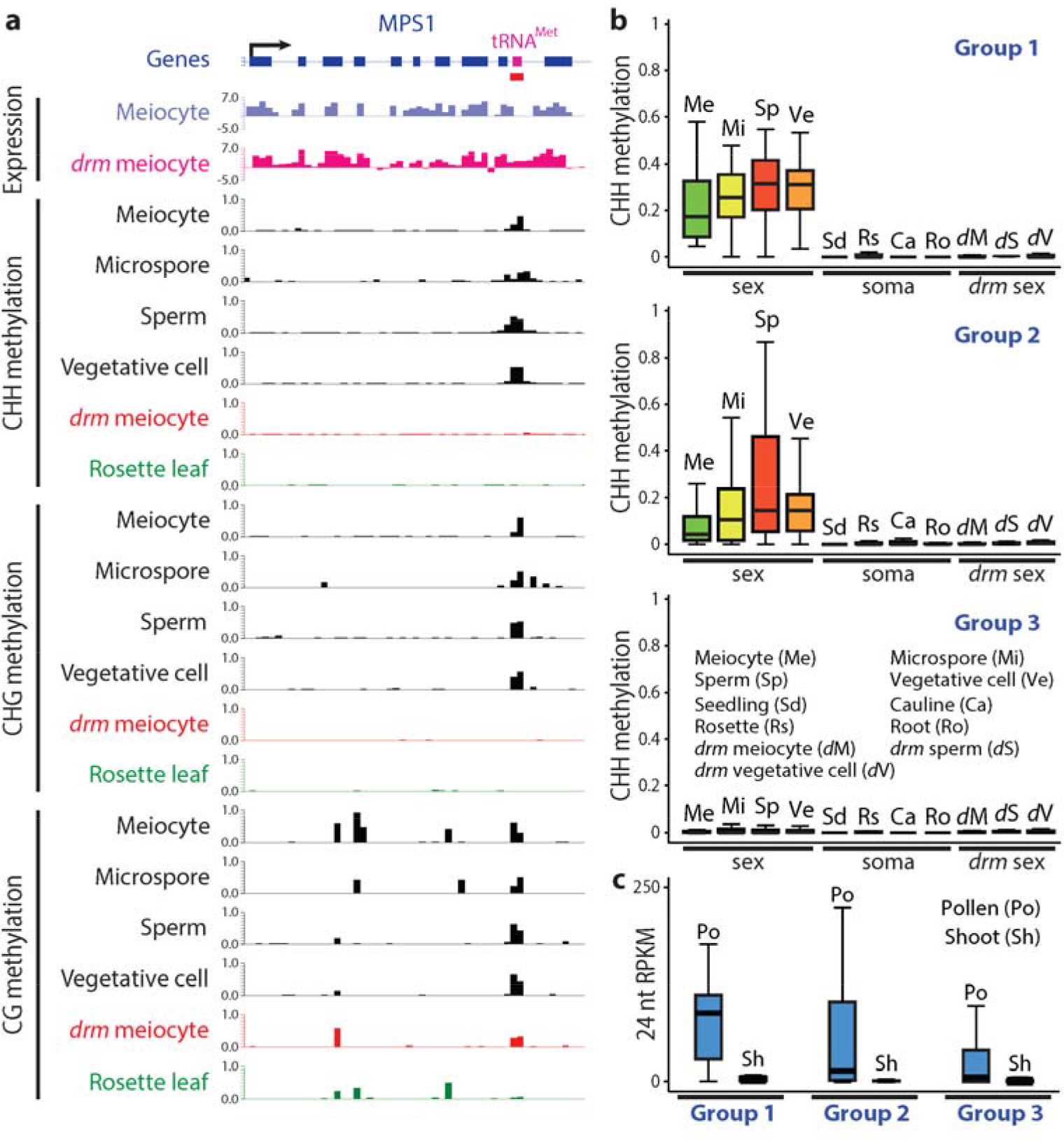
Pre-tRNA genes are hypermethylated in the male sexual lineage. **a**, Snapshots of transcription and DNA methylation (similar to Fig. 5b) at the methionine pre-tRNA locus (magenta box) in the last intron of the *MPS1* gene. *drm*, *drm1drm2*; SLM is underlined in red. Methylation patterns in other somatic tissues and *drm* sex cells are shown in Supplementary Fig. 6b. **b**, Box plots showing the absolute CHH methylation at three groups of pre-tRNA genes in sex cells (Me, meiocyte; Mi, microspore; Sp, sperm; Ve, vegetative cell), somatic tissues (Sd, seedling; Rs, rosette leaf; Ca, cauline leaf; Ro, root), and *drm* mutant sex cells (*d*M, *drm* meiocyte; *d*S, *drm* sperm; *d*V, *drm* vegetative cell). Group 1, the 24 pre-tRNA genes that overlap SLMs; Group 2, the additional 60 genes hypermethylated in the sexual lineage by RdDM; Group 3, the remaining 605 nuclear pre-tRNA genes. **c**, Box plots demonstrating the abundance of 24 nucleotide (nt) small RNA in pollen (Po) or shoot (Sh) at the abovementioned three groups of pre-tRNA genes.

### An SLM regulates the splicing of *MPS1* and is important for meiosis

One SLM-covered methionine pre-tRNA gene attracted our attention because it occurs within the last intron (between exons 9 and 10) of another gene called *MULTIPOLAR SPINDLE 1/PUTATIVE RECOMBINATION INITIATION DEFECTS 2* (*MPS1/PRD2*; Figs 6a and 7a, and Supplementary Fig. 6b). Given the emerging evidence of DNA methylation’s influence on splicing in plants^34,35^ and animals^36,37^, we examined our RNA-seq data to determine if the methylation status of this pre-tRNA locus affects *MPS1* splicing. Indeed, we detected cDNA reads that indicate incorrect splicing of *MPS1* at the last intron in *drm1drm2* mutant meiocytes (Fig. 6a). Quantitative RT-PCR analysis demonstrated that 28% of the mature *MPS1* mRNA retains the last intron in *drm1drm2* mutant meiocytes, whereas no such retention occurs in wild-type (Fig. 7a,b), confirming that the SLM within the intron is required for correct splicing of the *MPS1* transcript.

**Figure 7.**
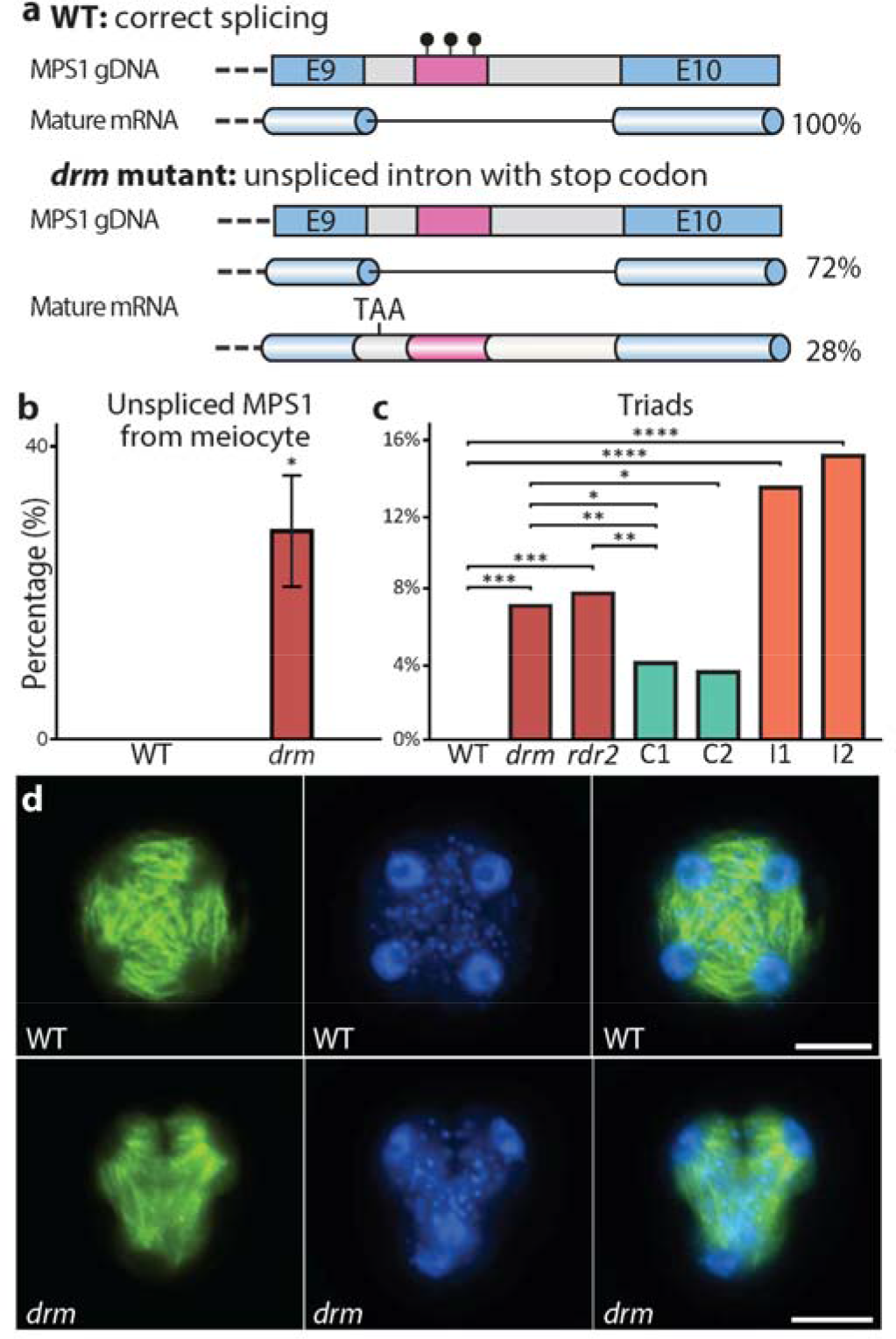
RdDM is important for the splicing of *MPS1* and normal meiosis. **a**, Gene model illustrating that the methionine pre-tRNA SLM (magenta bar) located in the last intron of *MPS1* affects the splicing of this intron. E, exon; black lollipops, DNA methylation. **b**, Quantitative RT-PCR showing the percentage of unspliced *MPS1* transcript in wild type (WT) and *drm1drm2* (*drm*) mutant meiocytes. \**P* < 0.02 (*t*-test). **c**, Percentage of meiotic triads in WT, *drm* and *rdr2* mutants, two complementation lines (C1 and C2) and two interference lines (I1 and I2) (\*\*\**P* < 1× 10^−7^, \*\**P* < 0.02, \**P* < 0.05, \*\*\*\**P* < 1× 10^−14^; Fisher’s exact test; C1, 514 observations; C2, 167 observations; for observation numbers of other genotypes refer to Supplementary Fig. 8 legend. **d**, Spindles (green) and nuclei (blue) of WT (tetrad) and *drm* (triad) meiotic products at tetrad stage. Scale bars, 10 μm.

We were intrigued by aberrant splicing of *MPS1* RNA in meiocytes because this gene is required for *Arabidopsis* meiosis^38,39^, and retention of the last intron introduces a premature stop codon (Fig. 7a). Furthermore, one of the described loss-of-function alleles affects splicing between exons 9 and 10, and another is an insertion in the intervening intron, indicating that exon 10 is essential for MPS1 activity^39^. We therefore analyzed meiotic progression in *drm1drm2* and *rdr2* mutants. Loss of MPS1 causes polyads – meiotic products numbering other than four^38^. Consistent with this phenotype, we found a significant occurrence of cellular triads in RdDM mutants (7.1% and 7.8% in *drm1drm2* and *rdr2*, respectively; Fig. 7c,d and Supplementary Fig. 8a-c). We also observed pentads in *drm1drm2* and *rdr2* mutants (Supplementary Fig. 8d,e), as has been reported for other RdDM mutants^40^, whereas we did not observe triads or pentads in wild type. Introduction of an *MPS1* transgene lacking the last intron into the *drm1drm2* background reduced the number of meiotic triads (4.1% and 3.6% for two independent complementation lines; Fig. 7c), but not to the undetectable level of wild-type plants. The persistence of triads suggested that the mis-spliced *MPS1* mRNA produces a protein that interferes with meiosis. To test this hypothesis, we introduced an *MPS1* transgene with mutations that prevent splicing of the last intron into wild-type plants (Supplementary Fig. 8f). The resulting transgenic plants exhibited a substantially higher percentage of meiotic triads than *drm1drm2* or *rdr2* mutants (13.4% and 15.1% for two independent interference lines; Fig. 7c and Supplementary Fig. 8g). Our results indicate that loss of methylation at the SLM within the last intron of *MPS1* causes intron retention and the production of an aberrant MPS1 protein that interferes with meiosis.

## DISCUSSION

Our results reveal the presence of a specific DNA methylation signature in the *Arabidopsis* male sexual lineage mediated by the RdDM pathway. SLMs suppress gene transcription and promote the splicing of a gene essential for meiosis, and are required for normal meiotic progression. This demonstrates that developmental gene regulation through DNA methylation reprogramming is not confined to gamete companion cells in flowering plants, and can occur through the deposition as well as the removal of methylation. Because RdDM appears to be ubiquitous in plant tissues, modulation of the RdDM pathway that achieves cell-specific methylation patterns can plausibly occur in any cell type. The epigenetic regulatory paradigm we describe here may therefore be broadly applicable to plant development.

SLMs are the product of developmentally orchestrated remodeling of DNA methylation via the RdDM pathway, but the small number of genes directly controlled by SLMs suggest that gene regulation is not the only, and perhaps not the main function of this remodeling. RdDM is known to target transposons^2,6,7^, there is a characterized tradeoff between methylation of transposons and gene expression^41^, and transposon suppression should be particularly important in cells that contribute to the next generation^26^. Therefore, RdDM may be balanced more aggressively in sex cells to ensure transposon silencing, even at the expense of gene expression – a phenomenon that might also occur in the shoot apical meristem, which gives rise to all above-ground plant cell types, including the gametes^42^. This would explain why we see sexual-lineage specific methylation, but very little if any soma-specific methylation. A more aggressive setting of the self *versus* non-self threshold in sex cells would also explain why sexual-lineage specific RdDM targets genes with such high frequency. Most SLMs may therefore be functionally neutral, or even slightly deleterious, and are likely evolutionarily transient, but a few, such as the one in *MPS1*, would be expected to confer a benefit and be retained through selection.

The substantial number (253) of SLMs that overlap genes in the *Arabidopsis* genome may elucidate a longstanding mystery regarding plant DNA methylation. The genes of flowering plants frequently exhibit CG-specific methylation of unclear origin and function^2^. This methylation has been hypothesized to arise due to transient RdDM activity^43^, which would have to occur in cells that contribute to the next generation – a description that fits SLMs. The remaining somatic CG methylation at SLMs (Fig. 4b and Supplementary Fig. 2d,e), which is maintained during somatic development without RdDM (Fig. 4c), provides evidence in support for this hypothesis. SLMs cover only a small fraction of the over 4000 genes with body methylation^44,45^, indicating that most body methylated genes are not presently targeted by RdDM in the male sexual lineage. However, shifting patterns of SLMs over thousands of generations could have plausibly created the existing gene body methylation pattern due to the strong trans-generational heritability of CG methylation^6^.

## METHODS

### Isolation of *A. thaliana* meiocytes, and sperm and vegetative cell nuclei

*A. thaliana* plants of Col-0 ecotype were grown under 16h light/ 8h dark in a growth chamber (21°C, 70% humidity). Stage 9 flower buds were collected and gently squeezed between a glass slide and coverslip. The released meiocytes (of meiotic prophase I) were examined carefully under a microscope. Clean meiocytes free from any somatic cell debris were transferred to a new slide using capillary glass pipettes, washed by 1 x PBS buffer 3 times, and frozen in liquid nitrogen. Sperm and vegetative cell nuclei were isolated as described previously^18^ (Supplementary Fig. 9).

### Sequencing library construction and analysis

Single-end bisulfite sequencing libraries for Illumina sequencing were constructed using the Ovation Ultralow Methyl-Seq Library Systems (Nugen, #0336) and EpiTect Fast Bisulfite Conversion (Qiagen, #59802) kits according to the kit protocols, except the incorporation of two rounds of bisulfite conversion. Bisulfite libraries were constructed from 2 biological replicates of wild-type (WT) meiocytes, *drm1*-*2drm2*-*2* (*drm1drm2*) mutant meiocytes, and *rdr2*-*1* mutant sperm and vegetative cell nuclei. Strand-specific RNA sequencing libraries were prepared using ScriptSeq v2 RNA-Seq (Illumina, #SSV21106) Library Preparation and Ovation RNA-Seq Systems for Model Organisms Arabidopsis (Nugen, #0351) kits, from 3 and 5 biological replicates of WT and *drm1drm2* mutant meiocytes, respectively, and 3 and 1 biological replicates of WT and *drm1drm2* mutant rosette leaf (from 40-day old plants). Bisulfite sequencing data from WT microspore^17^, sperm^18^, vegetative cell^18^, and embryo^18^, and *drm1drm2* mutant sperm and vegetative cell^28^ were used. Bisulfite sequencing data from 4 WT somatic tissues (in this section specifically referring to: cauline leaf^46^, rosette leaf^46,47^, roots^46,48^ and seedlings^48^), *rdd* mutant rosette leaf^29^, *drm2* and *rdr2* mutant seedlings^48^, and *drd1* mutant roots^48^ were obtained from published sources. Published sRNA data^49,50^, and bisulfite sequencing data from various root cell types^19^, and seedlings of *met1*-*1*, *pMET1::MET1 met1*-*1* complementation lines (T-MET1a and b) and their WT control plants^30^ were also used in this study.

Sequencing was performed at the DNA Sequencing Facility of the University of Cambridge Department of Biochemistry, BGI Tech Solutions Ltd. and the Vincent J. Coates Genomic Sequencing Laboratory at UC Berkeley. DNA methylation analysis was performed as previously described^18^, transcriptome analysis was performed using the Tophat-2.0.10 and Cufflinks-2.2.1 packages, sRNA abundances were calculated using Reads Per Kilobase per Million (RPKM; Fig. 3b) or Reads Per Million (RPM; Fig. 6c) of mapped 24-nt sRNA reads filtered for rRNAs.

### Transposon and gene meta analysis (ends analysis)

This was performed as described previously^18^.

### Identification of differentially methylated loci between the sexual lineage and somatic tissues

Fractional methylation in 50 bp windows across the genome was compared between an average of selected sex cells (SexAV) and an average of the four somatic tissues (SomAV) (Diff = SexAV - SomAV). CG and CHG methylation averages in sex cells were calculated using meiocytes, microspores and sperm, and CHH methylation average was calculated using microspores and sperm. We first selected windows meeting the following criterion: Diff_CG > 0 & Diff_CHG > 0 & Diff_CHH > 0 & (Diff_CG + Diff_CHG + Diff_CHH) > 0.3. The selected windows were merged to generate larger SLMs if they occurred within 100 bp. Merged SLMs were retained if they covered at least 100 bp, with significantly different levels of total methylation (Fisher’s exact test *P*-value < 0.001), having more methylation in all sex cell replicates than all somatic tissues, and met the following criterion: Diff_CG > 0 & Diff_CHG > 0.05 & Diff_CHH > 0.1 & (Diff_CG + Diff_CHG + Diff_CHH) > 0.4. This resulted in the identification of 1265 SLHs (used in Fig. 3a,b). The same criteria, except reversing the relationship between sexual lineage and somatic methylation, were used to identify 36 loci hypomethylated in the sexual lineage. SLHs were further refined by the criterion of having significantly (Fisher’s exact test *P*-value < 0.001) less methylation in sex cells (meiocyte, sperm and vegetative cell) of *drm1drm2* mutant in comparison to those of WT, leaving 1257 loci as a refined list of SLHs for further analyses (used in Fig. 3c). Refer to Supplementary Data 1 for abovementioned lists of loci.

The refined list of SLHs (1257 loci) was then separated into two groups based on the level of CHH/G methylation in somatic tissues: 1) SLMs with CHH and CHG methylation lower than 0.05 and 0.1, respectively, in all 4 somatic tissues (533 loci; Supplementary Data 1; used in Supplementary Fig. 3); 2) canonical SLHs with CHH methylation higher than 0.05 or CHG methylation higher than 0.1, in any of the 4 somatic tissues (724 loci; Supplementary Data 1; used in Fig. 4a). Both groups were analysed for overlap with published columella root cap DMRs (a merged list from reported C and CHH/G DMRs^19^): 173 and 469 of the canonical SLHs and SLMs have less than 10% overlap, respectively (Supplementary Data 1). The nonoverlapping 469 SLMs were used in subsequent analyses (Figs 4b-f, 5 and 6, and Supplementary Figs 4-7) as a refined list of SLMs.

### Identification of RdDM targets in sexual lineage and somatic tissues

Fractional methylation differences in 50 bp windows across the genome was compared between an average of RdDM mutants and WT tissues. For RdDM targets in sex cells, differential methylation was calculated using the average methylation in meiocyte, sperm and vegetative cells of WT subtracted by that of the *drm1drm2* mutant; for targets in somatic tissues, differential methylation was calculated using the average methylation in the 4 WT somatic tissues subtracted by the average methylation in *drm2* and *rdr2* mutant seedlings. For both targets, 50 bp windows meeting these criteria were selected: with a differential methylation larger than 0.05, 0.1 and 0.05 in CG, CHG and CHH contexts, respectively, and with a total differential methylation in CG, CHG and CHH contexts exceeding 0.2. Selected windows were subsequently merged to generate larger islands if they occurred within 100 bp. These islands were retained if they are at least 100 bp, with significantly different levels of total methylation (Fisher’s exact test *P*-value < 0.001), and meet the above-mentioned two criteria, except the 2^nd^ criterion uses a stricter cutoff of 0.3. This resulted in 9051 and 9993 RdDM targets in sex cells and somatic tissues, respectively.

### Box plots

All box plots follow this format: each box encloses the middle 50% of the distribution, with the horizontal line making the median, and vertical lines marking the minimum and maximum values that fall within 1.5 times the height of the box. Fig. 2b was generated using 50 bp windows with fractional CHH methylation larger than 0.3 in meiocytes compared to rosette leaf, and at least 20 informative sequenced cytosines in each of the 4 somatic tissues and 4 sex cells (meiocyte, microspore, sperm and vegetative cells). Fig. 3c was generated using 50 bp windows with substantial methylation in either WT sex cells or rosette leaf in the corresponding sequence context (cutoff settings: 0.3, 0.3 and 0.1 in CG, CHG and CHH contexts, respectively), and at least 10 informative sequenced cytosines in each replicate of the WT and *rdd* rosette leaf sample and 3 sex cells (meiocyte, microspore and sperm).

### RT-PCR

100 ng and 500 ng total RNA was reverse transcribed using RevertAid First Strand cDNA Synthesis Kit (Thermo Fisher scientific, #K1621) for quantitative RT-PCR (qRT-PCR; Figs 5c and 7b) and RT-PCR (Supplementary Fig. 8f), respectively. qRT-PCR was performed using SYBR Green (Roche, #4707516001) in triplicate on the LightCycler 480 Real-Time PCR System (Roche) and Cp values were averaged between 3 technical replicates to determine the target/reference ratio. Figs 5c and 7b show the averages of at least 3 biological replicates for each genotype or tissue type. RT-PCR was performed with 46 PCR cycles using primers JW134 and JW141 for *MPS1*, and 30 cycles using primers PHG34 and PHG101 for the control *ACTIN8*. *ACTIN8* was used as internal control in both RT- and qRT- PCRs, and all primers are listed in Supplementary Table 4.

### Meiosis microscopy

Slides were prepared from fixed *Arabidopsis* buds as previously described^51^. Cells were visualized and analyzed using a Nikon N*i*-E Eclipse fluorescent microscope (Nikon UK Ltd.) with NIS-elements imaging software. Triads were defined as cells with three nuclei that were observed without a boundary of overlap.

### Microtubule immunolocalization

Anthers were dissected from buds of the correct size (0.4-0.6mm long) and incubated in m-maleimidobenzoyl-N-hydroxysuccinimide ester (1mM in 1 х PBS and 0.05% [v/v] Triton X-100) for 30min^52^. Anthers were then fixed in 4% paraformaldehyde at 4°C for 1h. After one wash in 1 х PBS, anthers were digested in 0.5% (w/v) cytohelicase (Sigma), 0.3% (w/v) cellulase (Melford) and 0.3% (w/v) pectolyase (MP Biomedicals) in citrate buffer at 37°C for 75min. The enzyme mixture was then replaced with 1.5% Lipsol (Fisher Scientific) and transferred to Superfrost Plus slides (VWR). The digested anthers were spread with a mounted needle and then a coverslip was added. The slide was turned upside down and squashed onto the bench and then transferred to liquid nitrogen for 30s. The cover slip was immediately removed using a razor blade and the slide was allowed to air dry for 20min. An anti-α-Tubulin (Biorad) antibody (1:100 dilution in 1 х PBS and 1% BSA) was directly added to the slide and incubated overnight at 4°C. The slide was washed in 2 × 5min in 1 × PBS 0.1% Triton X-100 and then an anti-rat 488 DyLight (Vector labs) antibody (1:100 dilution in 1 × PBS and 1% BSA) was incubated for 30min at 37°C. The slide was then washed 2 × 5min in 1 × PBS 0.1% Triton X-100 and the excess buffer was allowed to drain off. DAPI in Vectashield (Vector labs) mounting medium was added to the slide.

### Construction of *MPS1* complementation and interference lines

To construct the *MPS1* complementation lines, overlapping PCR was carried out using primers PHG102, PHG103, PHG104 and PHG105 (Supplementary Table 4) to amplify the full *MPS1* genomic region covering the promoter, coding region and 3’ region with the exception of the last intron. To construct the *MPS1* interference lines with splicing site mutations, DNA was amplified by PCR for the full *MPS1* genomic region, using primers PHG102 and PHG103. Two point mutations at the splicing sites of the last intron were created using the QuikChange Lightning Multi Site-Directed Mutagenesis Kit (Agilent, #210515) using primers PHG106 and PHG107. Both entry clones were sequenced and cloned into destination vectors *pGWB13*-*Bar* (a modified *pGWB13*^53^ vector with BASTA resistance gene replacing Hygromycin and Kanamycin resistance genes) and *pMDC107*-*NTF* (a modified pMDC107 vector with NTF^54^ replacing mGFP6) via Gateway cloning, and transformed into *drm1drm2* mutant and wild-type plants to generate the complementation lines (*pGWB13* for C1 and *pMDC107* for C2) and interference lines (*pGWB13* for I1 and *pMDC107* for I2). 5, 2, 5 and 4 T1 generation plants of the C1, C2, I1 and I2 lines, respectively, were used in meiotic phenotype analysis.

### Data availability

All sequencing data that support the findings of this study have been deposited in the National Center for Biotechnology Information Gene Expression Omnibus (GEO) under accession number GSE86583. All other relevant data are available from the corresponding author on request.

## ACKNOWLEDGMENTS

We thank Daniel Zilberman for intellectual contributions to this work, and Daniel Zilberman, Caroline Dean, Kirsten Bomblies, Vinod Kumar, Siobhan Brady and Sophien Kamoun for comments on the manuscript, Hugh Dickinson and Josephine Hellberg for developing the meiocyte isolation method, Giles Oldroyd for the *pGWB13*-*Bar* vector, Elisa Fiume for the *pMDC107*-*NTF* vector, Matthew Hartley, Matthew Couchman and Tjelvar Sten Gunnar Olsson for bioinformatics support, and the John Innes Centre Bioimaging Facility for their assistance with microscopy. This work was funded by a Biotechnology and Biological Sciences Research Council (BBSRC) David Phillips Fellowship (BBL0250431) to X.F., a BBSRC grant (BBM01973X1) to J.H., and a Sainsbury PhD Studentship to J.W.

## AUTHOR CONTRIBUTIONS

X.F. designed the study, J.W., H.G., J.Z., B.A., K.F., J.H. and X.F. performed the experiments, J.W., H.G., M.V. and X.F. analyzed the data, X.F. wrote the manuscript.

## AUTHOR INFORMATION

The authors declare no competing financial interests. Correspondence and requests for materials should be addressed to X.F.

---

**Supplementary Figure 1:**
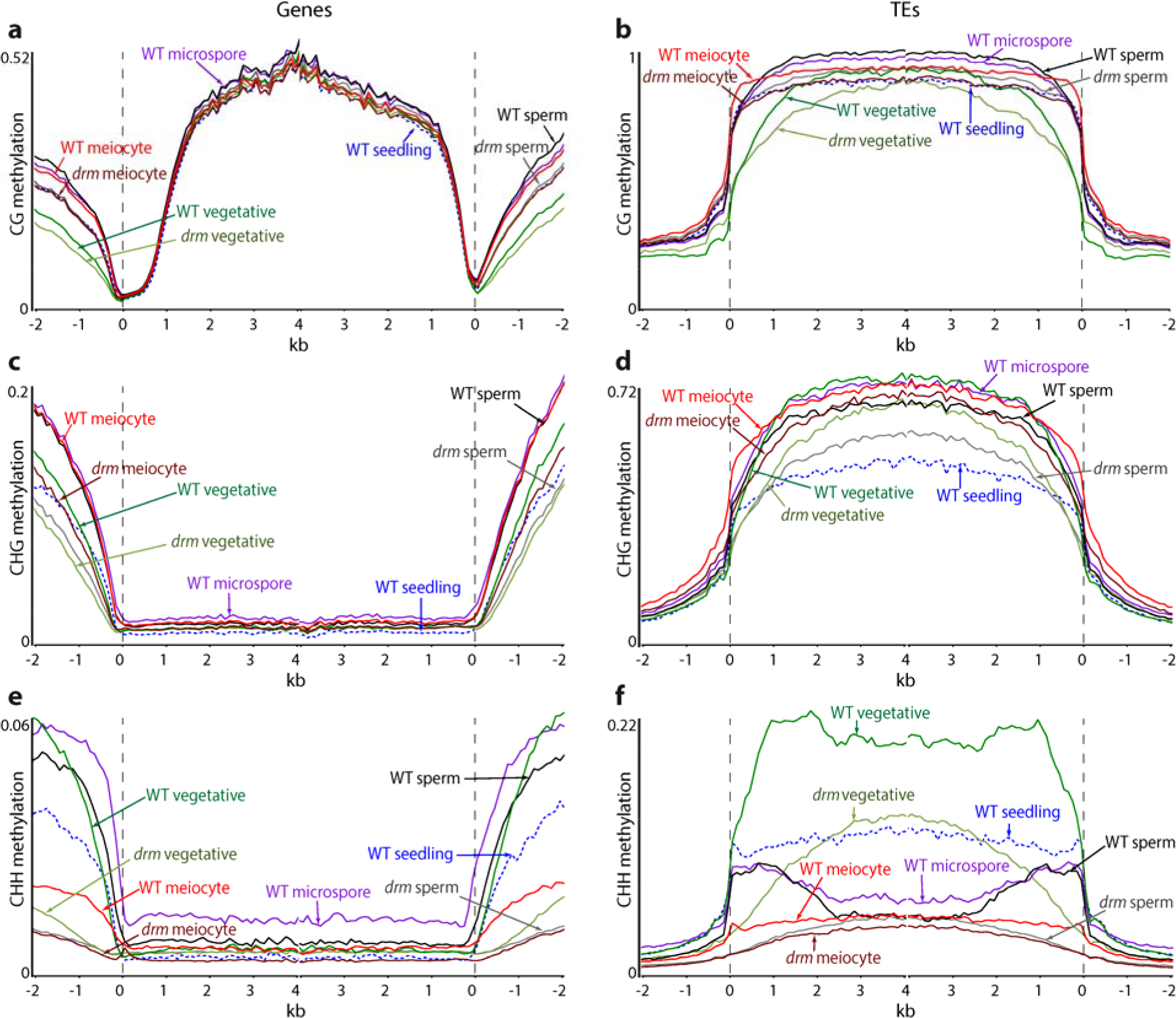
DNA methylation profiles at genes and transposons in *Arabidopsis* male sexual lineage cells. *A. thaliana* genes (a, c, e) or transposable elements (TEs; b, d, f) were aligned at the 5’ end (left panel) or the 3’ end (right panel), and average methylation levels in the CG (a-b), CHG (c-d) or CHH (e-f) context for each 100-bp interval are plotted. *drm*, *drm1drm2*. The dashed line at zero represents the point of alignment.

**Supplementary Figure 2:**
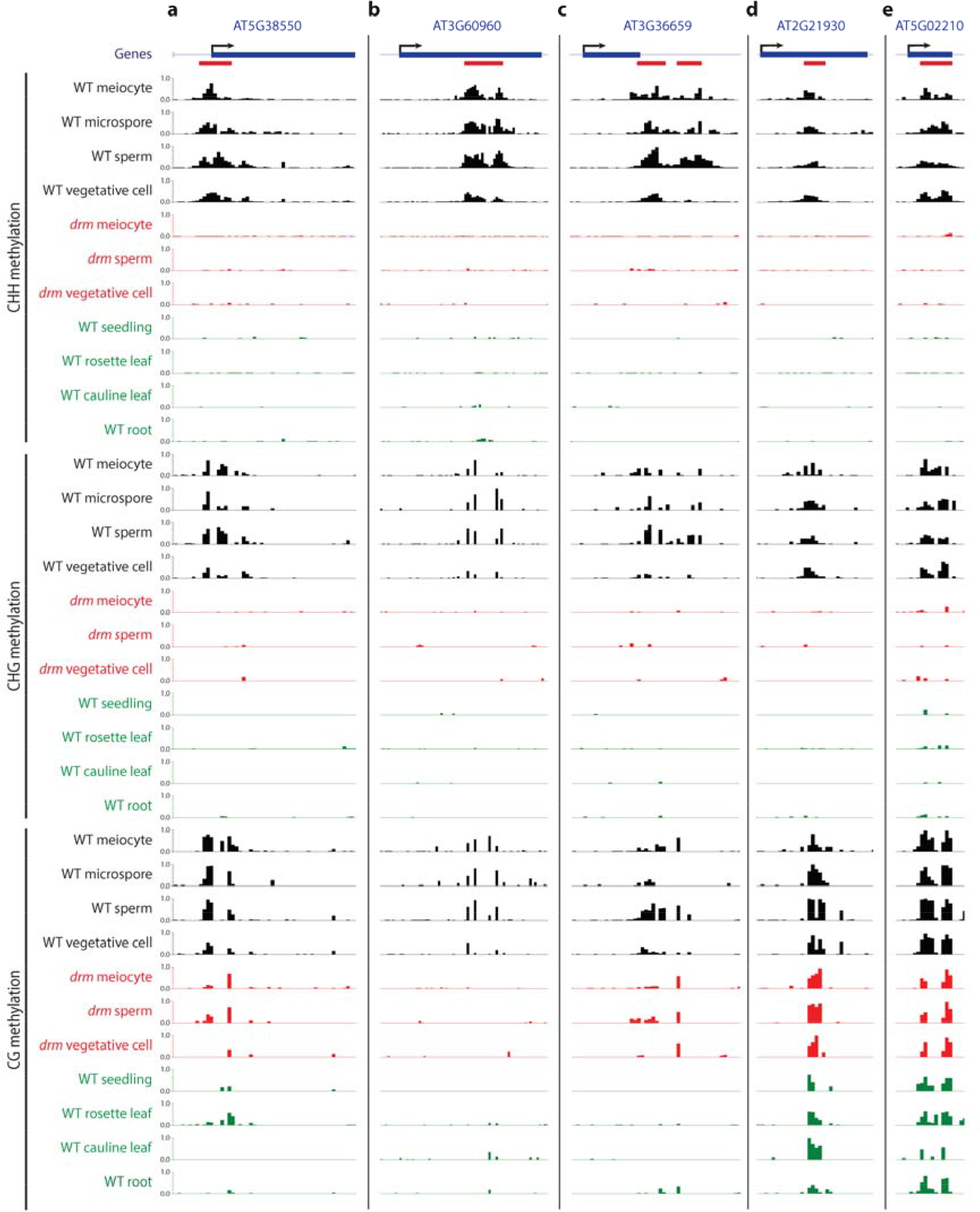
Examples of sexual-lineage hypermethylated loci (SLHs). Examples of typical SLHs located at the transcriptional start and termination sites and body of genes (a-c), and SLHs (that are also SLMs) with remnant CG methylation in *drm1drm2* (*drm*) mutant sex cells and wild-type somatic tissues (d, e). SLHs are underlined in red (refer to Supplementary Data 1 for a full list).

**Supplementary Figure 3:**
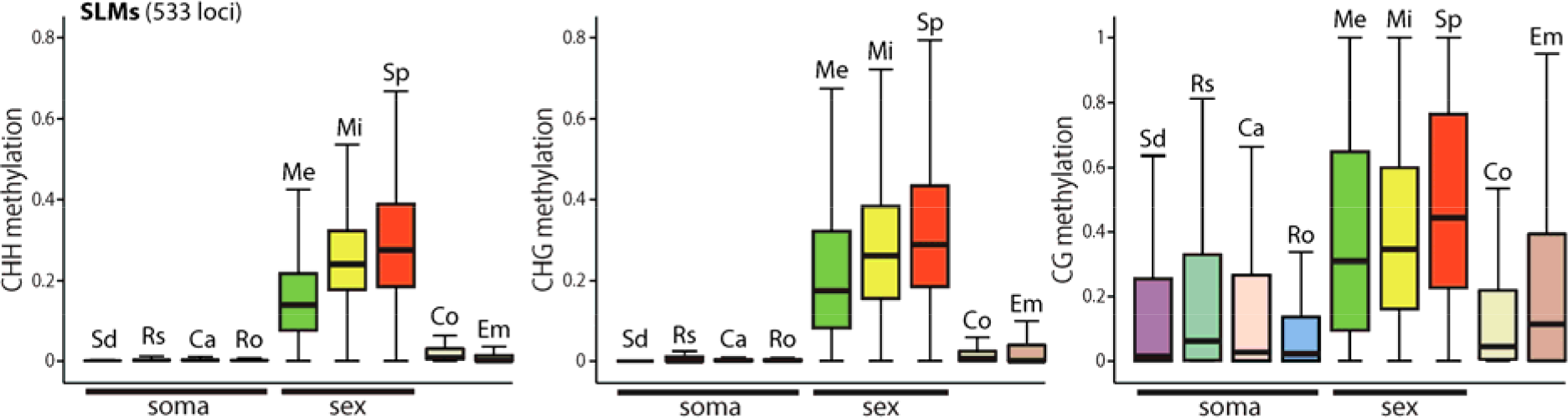
SLMs have little CHH/G methylation in columella and embryo. Box plots showing the absolute methylation at SLMs (533 loci before filtering out columella overlaps; see Methods) in somatic tissues (Sd, seedling; Rs, rosette leaf; Ca, cauline leaf; Ro, root), sex cells (Me, meiocyte; Mi, microspore; Sp, sperm), columella root cap (Co) and embryo (Em).

**Supplementary Figure 4:**
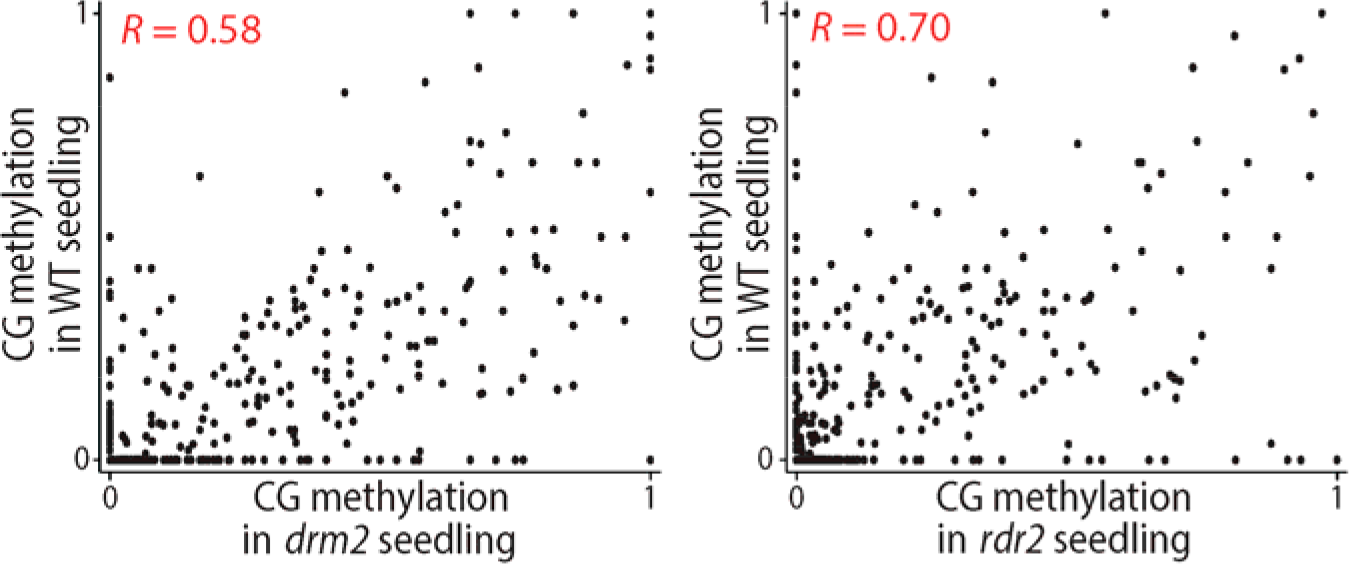
CG methylation in seedlings of wild type (WT) and RdDM mutants is strongly correlated. Scatter plots showing linear correlation between CG methylation at SLMs in seedlings of WT and *drm2* (Pearson’s *R* = 0.58), and WT and *rdr2* (Pearson’s *R* = 0.70).

**Supplementary Figure 5:**
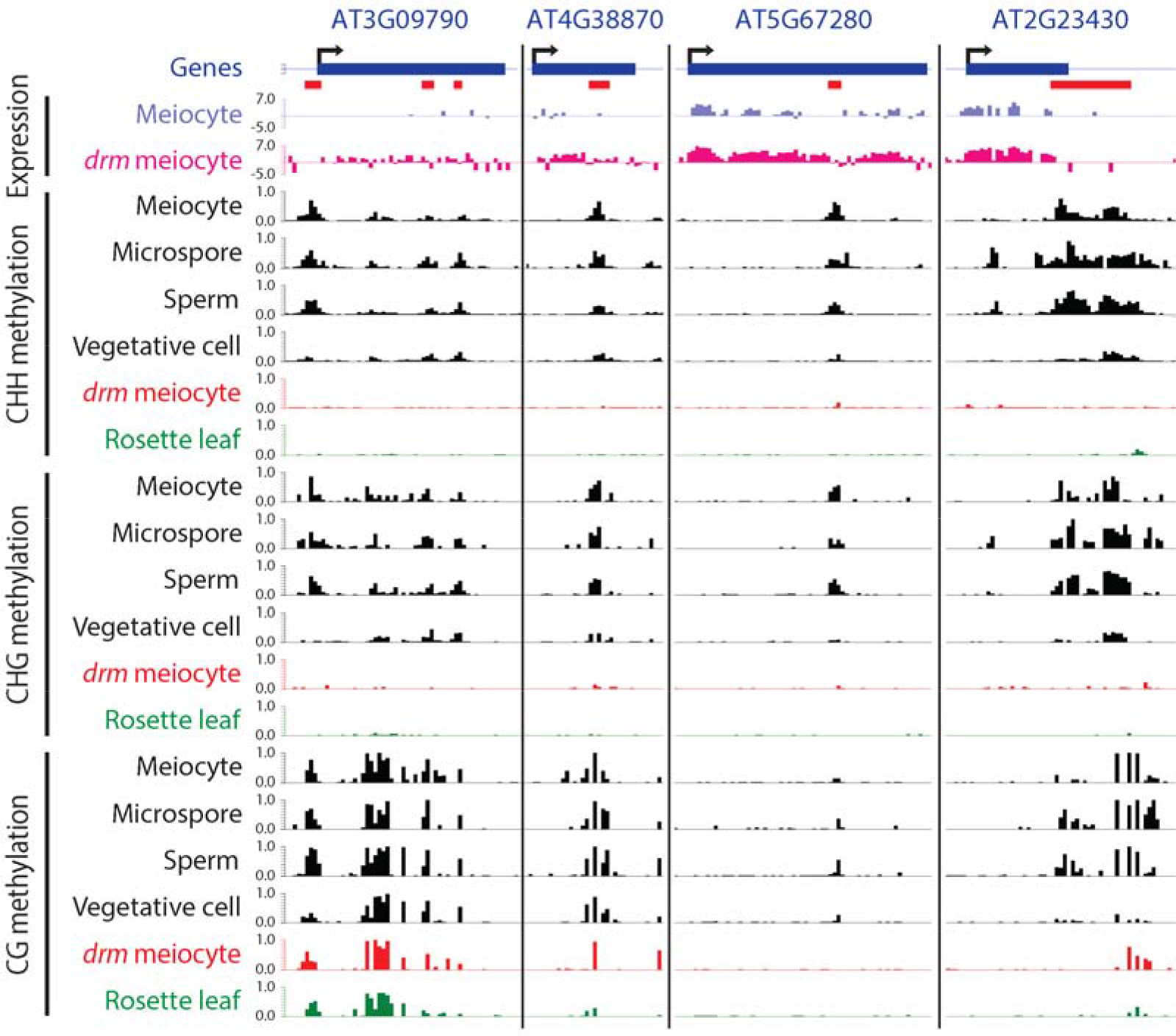
Examples of genes suppressed by sexual-lineage-specific methylation in meiocytes. Similar to Fig. 5b, snapshots of cytosine methylation in wild-type male sex cells, *drm1drm2* (*drm*) meiocyte, and wild-type rosette leaves, and transcriptional expression (in log_2_RPKM) in wild-type and *drm* meiocyte are shown. SLMs are underlined in red.

**Supplementary Figure 6:**
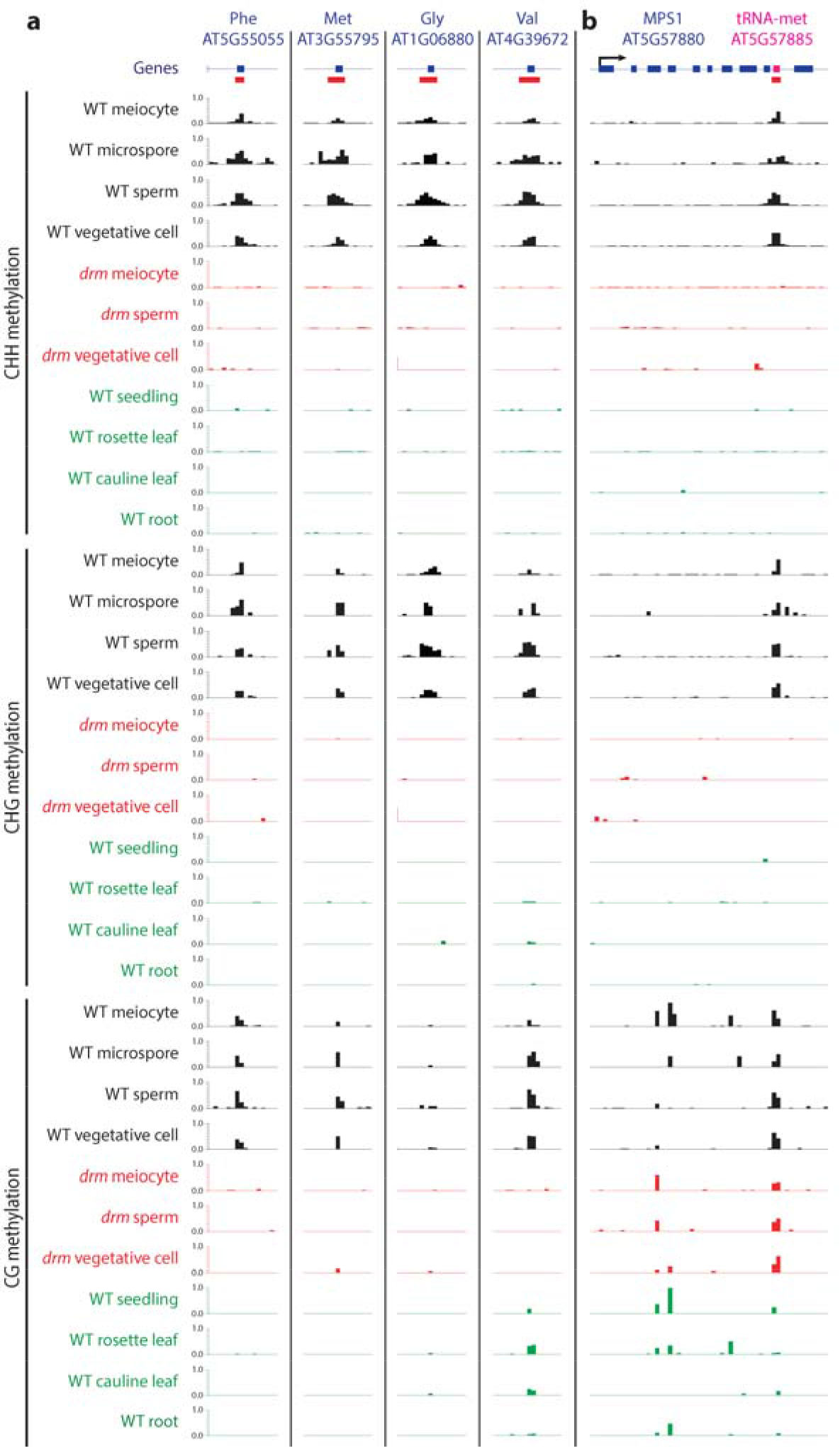
Examples of sexual-lineage-specific methylation at pre-tRNA genes. Snapshots of cytosine methylation, similar to Supplementary Fig. 2, in wild-type (WT) male sex cells (black), *drm1drm2* (*drm*) mutant sex cells (red), and four somatic tissues (green). SLMs are underlined in red. **a**, Examples of SLMs at pre-tRNA genes encoding phenylalanine, methionine, glycine or valine anticodons. **b**, SLM at the methionine pre-tRNA gene (magenta box) located in the last intron of *MPS1*.

**Supplementary Figure 7.**
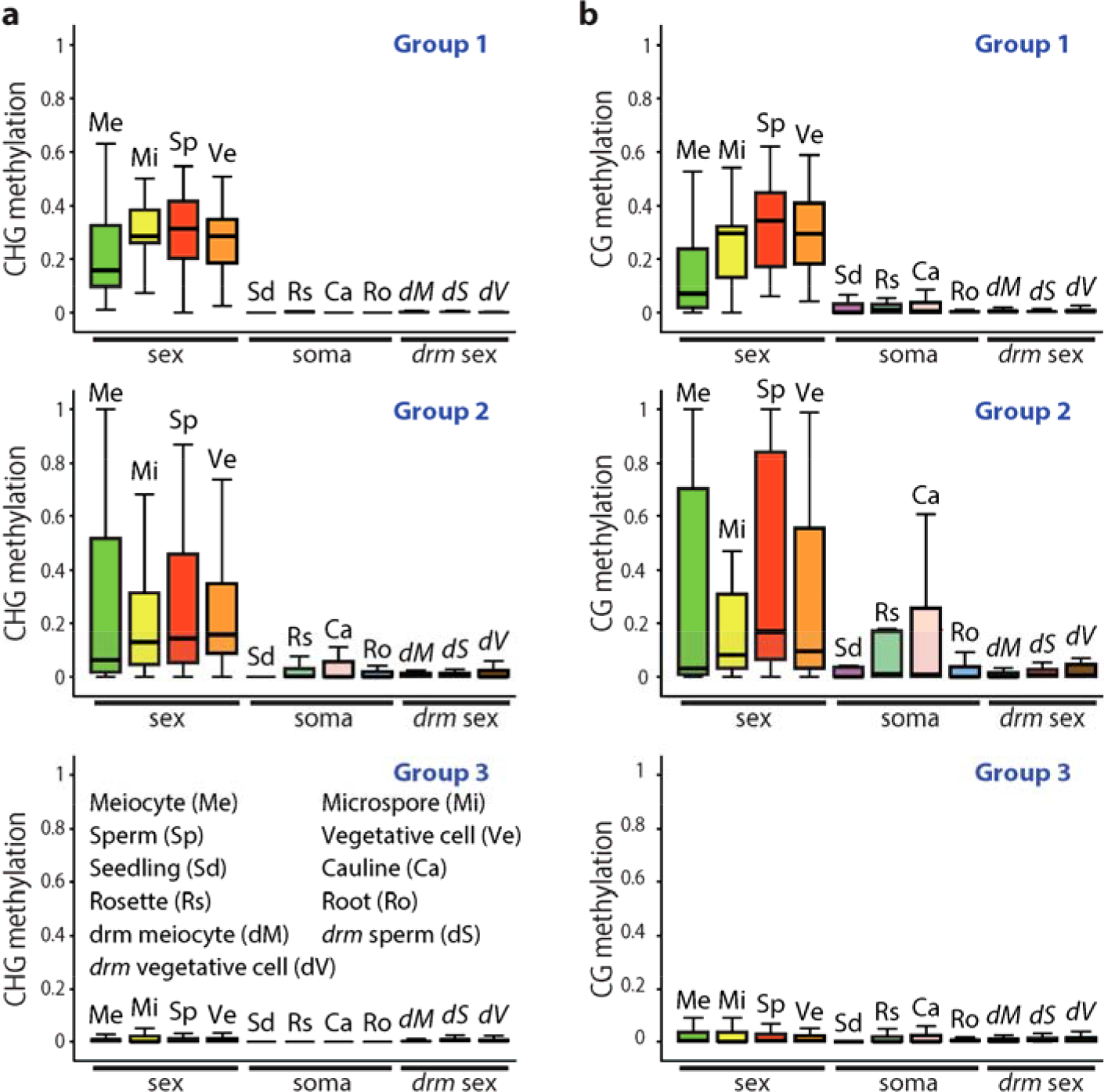
Pre-tRNA genes are hypermethylated in male sex cells. Similar to Fig. 6b, these box plots show the absolute CHG (a) and CG (b) methylation at 3 groups of pre-tRNA genes in sex cells (Me, meiocyte; Mi, microspore; Sp, sperm; Ve, vegetative cell), somatic tissues (Sd, seedling; Rs, rosette leaf; Ca, cauline leaf; Ro, root), and *drm* (*drm1drm2*) mutant sex cells (*d*M, *drm* meiocyte; *d*S, *drm* sperm; *d*V, *drm* vegetative cell). Refer to Fig. 6 legend for the 3 groups of pre-tRNA genes.

**Supplementary Figure 8:**
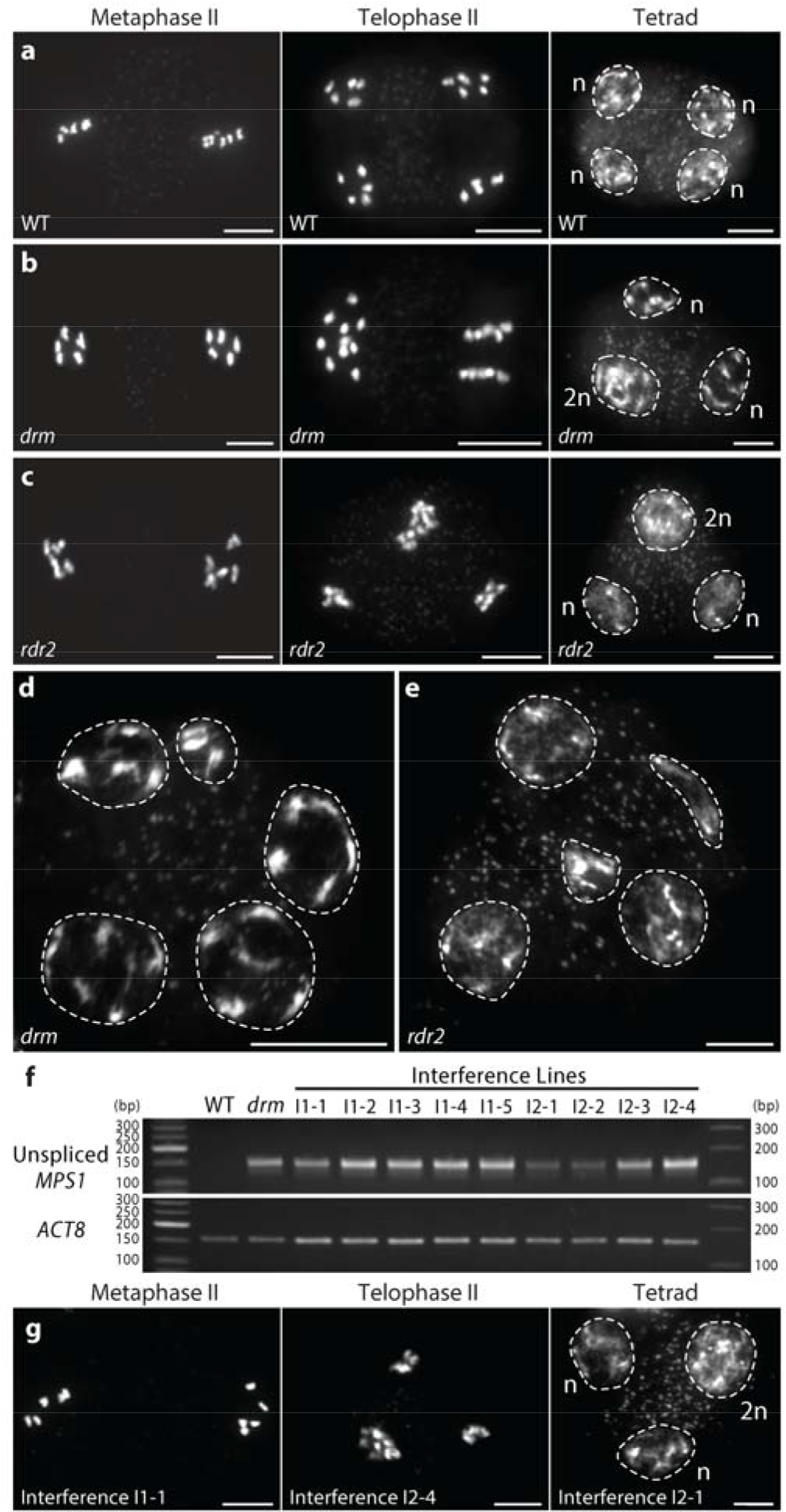
Meiotic defects in RdDM mutants and *MPS1* interference lines. **a-e**, Male meiosis II in wild type (WT; a), *drm1drm2* (*drm*; b, d) and *rdr2* (c, e) mutants, and the *MPS1* interference lines (g). All instances of WT male meiosis we observed (301 observations) were normal and lead to tetrads at the end of meiosis II (a). However, in 7.1% (380 total observations) and 7.8% (502 total observations) instances of *drm* (b) and *rdr2* (c) male meiosis, respectively, chromosomes fail to separate, so that triads are observed at telophase II. Occasionally we also observed pentads in *drm* (d) and *rdr2* (e) mutants. **f**, RT-PCR showing the expression of *MPS1* transcript retaining last intron (149 bp) in *drm* mutant and 9 T1 plants of the interference lines, but not in WT. *ACT8* as control shows 156 bp bands. **g**, Interference lines exhibit even higher percentages of triads (I1, 13.4%, 463 total observations; I2, 15.1%, 292 total observations). n, the number of chromosomes in the haploid genome. Scale bars, 10 μm.

**Supplementary Figure 9:**
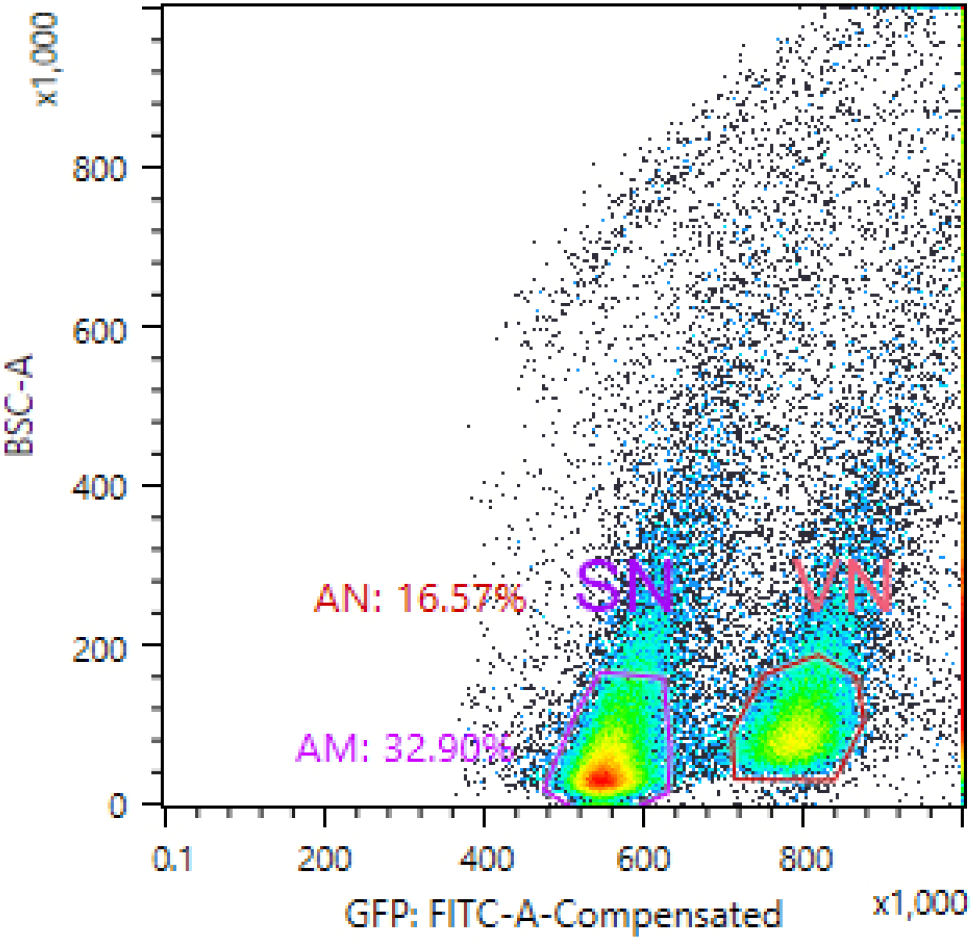
Separation of sperm and vegetative cell nuclei via fluorescence-activated cell sorting (FACS). A representative flow cytometry plot showing two clear populations which indicate SYBR Green-stained sperm nuclei (SN) and vegetative cell nuclei (VN), respectively. AM and AN indicate the percentages of sorted SN and VN in total events, respectively.

**Supplementary Table 1:** Sequencing summary statistics for bisulfite sequencing libraries. Mean DNA methylation (Met) was calculated by averaging methylation of individual cytosines in each context, and chloroplast CHH methylation was used as a measure of cytosine non-conversion and other errors.

| **Sample** | **Read Length** | **Number of Reads** | **Nuclear Genome Coverage** | **Nuclear CG Met** | **Nuclear CHG Met** | **Nuclear CHH Met** | **Chloroplast CHH Met** |
| --- | --- | --- | --- | --- | --- | --- | --- |
| **Wild-type Meocyte Rep1** | 100 | 219966493 | 104.2 | 28.8% | 13.3% | 1.5% | 0.4% |
| **Wild-type Meocyte Rep2** | 150 | 34237362 | 19.1 | 27.8% | 13.0% | 1.2% | 0.3% |
| ***drm1drm2* Meocyte Rep1** | 75 | 83452411 | 30.8 | 24.4% | 9.9% | 0.9% | 0.3% |
| ***drm1drm2* Meocyte Rep2** | 75 | 81120807 | 29.9 | 26.7% | 11.7% | 0.9% | 0.2% |
| ***rdr2* Sperm** | 100 | 239282250 | 132.7 | 29.2% | 10.2% | 1.2% | 0.6% |
| ***rdr2* Vegetative Cell** | 100 | 41176080 | 26.1 | 25.6% | 11.5% | 2.7% | 0.7% |

**Supplementary Table 2:** The percentages of RdDM targets overlapping, within 500 bp, and more than 500 bp from genes or transposons. Two methods were used to select sets of loci that mimic SLMs (or canonical SLHs) in relation to genes for (b). For SLMs: Method 1 randomly selected 469 loci of the same size distribution as SLMs, and with 253 of them overlapping gene body, 93 within 500 bp of genes, and 123 more than 500 bp from genes; in Method 2, we randomly selected 253 loci overlapping genes and of the same sizes as the 253 gene-overlapping SLMs, and did similarly for the 93 and 123 loci that are less and more than 500 bp from genes, respectively, for a total of 469 loci. For canonical SLHs, the same approach was used to generate two corresponding lists that mimic canonical SLHs in relation to genes.

**a**
|  | Number of loci overlapping gene body | Percentage of loci overlapping gene body | Number of loci overlapping 500bp 5' or 3' of genes | Percentage of loci overlapping 500bp 5' or 3' of genes | Number of loci away from genes for more than 500 bp | Percentage of loci away from genes for more than 500 bp | Total number of loci |
| --- | --- | --- | --- | --- | --- | --- | --- |
| **SLMs** | 253 | 54.0% | 93 | 19.8% | 123 | 26.2% | 469 |
| **Canonical SLHs** | 169 | 23.3% | 209 | 28.9% | 346 | 47.8% | 724 |
| **Sex cell RdDM targets** | 717 | 9.3% | 2029 | 26.4% | 4950 | 64.3% | 7696 |
| **Soma RdDM targets** | 794 | 7.9% | 2380 | 23.8% | 6819 | 68.2% | 9993 |

**b**
|  | Number of loci overlapping TEs | Percentage of loci overlapping TEs | Number of loci overlapping 500bp 5' or 3' of TEs | Percentage of loci overlapping 500bp 5' or 3' of TEs | Number of loci away from TEs for more than 500 bp | Percentage of loci away from TEs for more than 500 bp | Total number of loci |
| --- | --- | --- | --- | --- | --- | --- | --- |
| **SLMs** | 84 | 17.9% | 75 | 16.0% | 310 | 66.1% | 469 |
| **Canonical SLHs (cSLHs)** | 470 | 64.9% | 67 | 9.3% | 187 | 25.8% | 724 |
| **Sex cell RdDM targets** | 6039 | 78.5% | 525 | 6.8% | 1132 | 14.7% | 7696 |
| **Soma RdDM targets** | 7794 | 78.0% | 797 | 8.0% | 1402 | 14.0% | 9993 |
| **Randomly selected SLM-like loci (Method1)** | 99 | 21.1% | 52 | 11.1% | 318 | 67.8% | 469 |
| **Randomly**<br>**selected**<br>**SLM-like**<br>**loci**<br>**(Method2)** | 103 | 22.0% | 47 | 10.0% | 319 | 68.0% | 469 |
| **Randomly**<br>**selected**<br>**cSLH-like**<br>**loci**<br>**(Method1)** | 301 | 41.6% | 68 | 9.4% | 355 | 49.0% | 724 |
| **Randomly**<br>**selected**<br>**cSLH-like**<br>**loci**<br>**(Method2)** | 293 | 40.5% | 74 | 10.2% | 357 | 49.3% | 724 |

**Supplementary Table 3:** pre-tRNA genes hypermethylated by RdDM in the sexual lineage. Tables list the anticodons of the 24 pre-tRNA genes overlapping SLMs (a) and a complete list of 84 pre-tRNA genes methylated in the sexual lineage by RdDM (b).

| Amino acid | Anticodon | Number of pre-tRNA genes covered by SLMs | Number of nuclear pre-tRNA genes per anticodon | Percentage of nuclear pre-tRNA genes covered by SLMs |
| --- | --- | --- | --- | --- |
| **Phe** | GAA | 12 | 16 | 75.0% |
| **Val** | CAC | 2 | 8 | 25.0% |
| **Met** | CAT | 5 | 24 | 20.8% |
| **Cys** | GCA | 3 | 15 | 20.0% |
| **Ala** | CGC | 1 | 7 | 14.3% |
| **Gly** | GCC | 1 | 23 | 4.3% |
| **TOTAL** |  | 24 | 93 | 25.8% |

**b**
| Amino acid | Anticodon | Number of pre-tRNA genes hypermethylated by RdDM in the sexual lineage | Number of nuclear pre-tRNA genes per anticodon | Percentage of nuclear pre-tRNA genes hypermethylated by RdDM in the sexual lineage |
| --- | --- | --- | --- | --- |
| **Phe** | GAA | 16 | 16 | 100.0% |
| **Gly** | ACC | 1 | 1 | 100.0% |
| **Trp** | CCA | 12 | 14 | 85.7% |
| **Val** | CAC | 6 | 8 | 75.0% |
| **Cys** | GCA | 11 | 15 | 73.3% |
| **Ile** | TAT | 3 | 5 | 60.0% |
| **Gly** | GCC | 13 | 23 | 56.5% |
| **Met** | CAT | 10 | 24 | 41.7% |
| **Ala** | CGC | 1 | 7 | 14.3% |
| **Ser** | AGA | 4 | 37 | 10.8% |
| **Leu** | TAG | 1 | 10 | 10.0% |
| **Tyr** | GTA | 6 | 76 | 7.9% |
| **TOTAL** |  | 84 | 236 | 35.6% |

**Supplementary Table 4:**
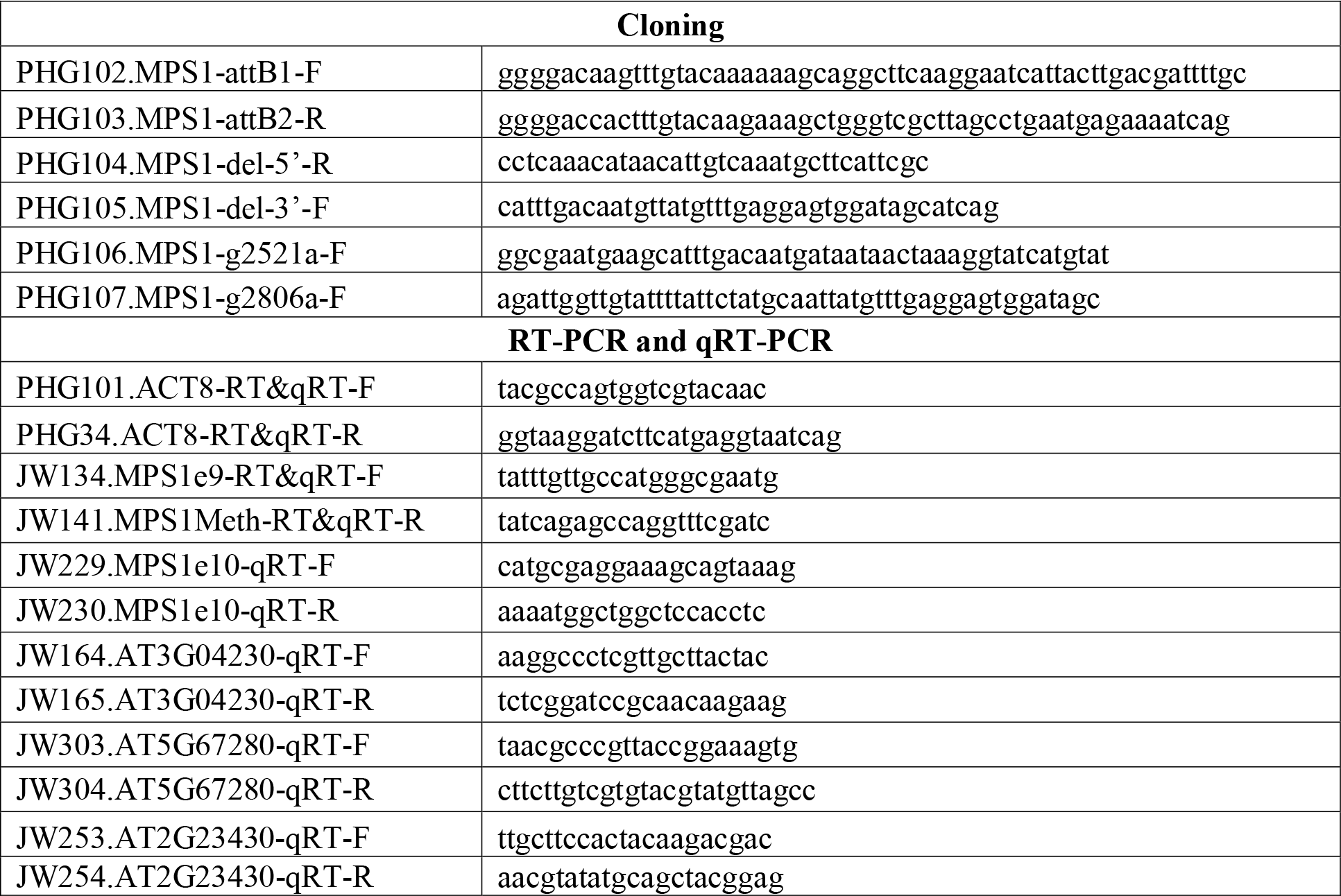
Primers used in this study.

**Supplementary Data Materials for this manuscript include the following:**

**Supplementary Data 1:** List of loci that are significantly differentially methylated between the sexual lineage and somatic tissues

**Supplementary Data 2:** Expression of 83 meiotic genes in our meiocyte RNA-seq data in comparison to those from published meiocyte transcriptomes

**Supplementary Data 3:** List of differentially expressed genes and their association with SLMs

## REFERENCES

1 Zemach, A. & Zilberman, D. Evolution of eukaryotic DNA methylation and the pursuit of safer sex. Curr Biol 20, R780–785, doi:10.1016/j.cub.2010.07.007 (2010).

2 Law, J. A. & Jacobsen, S. E. Establishing, maintaining and modifying DNA methylation patterns in plants and animals. Nat Rev Genet 11, 204–220, doi:10.1038/nrg2719 (2010).

3 He, X. J., Chen, T. & Zhu, J. K. Regulation and function of DNA methylation in plants and animals. Cell Res 21, 442–465, doi:10.1038/cr.2011.23 (2011).

4 Du, J., Johnson, L. M., Jacobsen, S. E. & Patel, D. J. DNA methylation pathways and their crosstalk with histone methylation. Nat Rev Mol Cell Biol 16, 519–532, doi:10.1038/nrm4043 (2015).

5 Pikaard, C. S. & Mittelsten Scheid, O. Epigenetic regulation in plants. Cold Spring Harb Perspect Biol 6, a019315, doi:10.1101/cshperspect.a019315 (2014).

6 Matzke, M. A. & Mosher, R. A. RNA-directed DNA methylation: an epigenetic pathway of increasing complexity. Nat Rev Genet 15, 394–408, doi:10.1038/nrg3683 (2014).

7 Kim, M. Y. & Zilberman, D. DNA methylation as a system of plant genomic immunity. Trends Plant Sci 19, 320–326, doi:10.1016/j.tplants.2014.01.014 (2014).

8 Lipka, D. B. et al. Identification of DNA methylation changes at cis-regulatory elements during early steps of HSC differentiation using tagmentation-based whole genome bisulfite sequencing. Cell Cycle 13, 3476–3487, doi:10.4161/15384101.2014.973334 (2014).

9 Dawlaty, M. M. et al. Loss of Tet enzymes compromises proper differentiation of embryonic stem cells. Dev Cell 29, 102–111, doi:10.1016/j.devcel.2014.03.003 (2014).

10 Kubo, N. et al. DNA methylation and gene expression dynamics during spermatogonial stem cell differentiation in the early postnatal mouse testis. BMC Genomics 16, 624, doi:10.1186/s12864-015-1833-5 (2015).

11 Slieker, R. C. et al. DNA Methylation Landscapes of Human Fetal Development. Plos Genet 11, e1005583, doi:10.1371/journal.pgen.1005583 (2015).

12 Xie, W. et al. Epigenomic analysis of multilineage differentiation of human embryonic stem cells. Cell 153, 1134–1148, doi:10.1016/j.cell.2013.04.022 (2013).

13 Cacchiarelli, D. et al. Integrative Analyses of Human Reprogramming Reveal Dynamic Nature of Induced Pluripotency. Cell 162, 412–424, doi:10.1016/j.cell.2015.06.016 (2015).

14 Benner, C., Isoda, T. & Murre, C. New roles for DNA cytosine modification, eRNA, anchors, and superanchors in developing B cell progenitors. Proc Natl Acad Sci U S A 112, 12776–12781, doi:10.1073/pnas.1512995112 (2015).

15 Rodrigues, J. A. & Zilberman, D. Evolution and function of genomic imprinting in plants. Genes Dev 29, 2517–2531, doi:10.1101/gad.269902.115 (2015).

16 Park, K. et al. DNA demethylation is initiated in the central cells of Arabidopsis and rice. Proc Natl Acad Sci U S A 113, 15138–15143, doi:10.1073/pnas.1619047114 (2016).

17 Calarco, J. P. et al. Reprogramming of DNA methylation in pollen guides epigenetic inheritance via small RNA. Cell 151, 194–205, doi:10.1016/j.cell.2012.09.001 (2012).

18 Ibarra, C. A. et al. Active DNA demethylation in plant companion cells reinforces transposon methylation in gametes. Science 337, 1360–1364, doi:10.1126/science.1224839 (2012).

19 Kawakatsu, T. et al. Unique cell-type-specific patterns of DNA methylation in the root meristem. Nat Plants 2, 16058, doi:10.1038/nplants.2016.58 (2016).

20 Hsieh, T. F. et al. Genome-wide demethylation of Arabidopsis endosperm. Science 324, 1451–1454, doi:10.1126/science.1172417 (2009).

21 Secco, D. et al. Stress induced gene expression drives transient DNA methylation changes at adjacent repetitive elements. Elife 4, doi:10.7554/eLife.09343 (2015).

22 Jiang, C. et al. Environmentally responsive genome-wide accumulation of de novo Arabidopsis thaliana mutations and epimutations. Genome Res 24, 1821–1829, doi:10.1101/gr.177659.114 (2014).

23 Dowen, R. H. et al. Widespread dynamic DNA methylation in response to biotic stress. Proc Natl Acad Sci U S A 109, E2183–2191, doi:10.1073/pnas.1209329109 (2012).

24 Bilichak, A., Ilnystkyy, Y., Hollunder, J. & Kovalchuk, I. The progeny of Arabidopsis thaliana plants exposed to salt exhibit changes in DNA methylation, histone modifications and gene expression. PLoS One 7, e30515, doi:10.1371/journal.pone.0030515 (2012).

25 Wibowo, A. et al. Hyperosmotic stress memory in Arabidopsis is mediated by distinct epigenetically labile sites in the genome and is restricted in the male germline by DNA glycosylase activity. Elife 5, doi:10.7554/eLife.13546 (2016).

26 Feng, X., Zilberman, D. & Dickinson, H. A conversation across generations: soma-germ cell crosstalk in plants. Dev Cell 24, 215–225, doi:10.1016/j.devcel.2013.01.014 (2013).

27 Kawashima, T. & Berger, F. Epigenetic reprogramming in plant sexual reproduction. Nat Rev Genet 15, 613–624, doi:10.1038/nrg3685 (2014).

28 Hsieh, P. H. et al. Arabidopsis male sexual lineage exhibits more robust maintenance of CG methylation than somatic tissues. Proc Natl Acad Sci U S A 113, 15132–15137, doi:10.1073/pnas.1619074114 (2016).

29 Stroud, H., Greenberg, M. V., Feng, S., Bernatavichute, Y. V. & Jacobsen, S. E. Comprehensive analysis of silencing mutants reveals complex regulation of the Arabidopsis methylome. Cell 152, 352–364, doi:10.1016/j.cell.2012.10.054 (2013).

30 Catoni, M. et al. DNA sequence properties that predict susceptibility to epiallelic switching. Embo J 36, 617–628, doi:10.15252/embj.201695602 (2017).

31 Chen, C. et al. Meiosis-specific gene discovery in plants: RNA-Seq applied to isolated Arabidopsis male meiocytes. BMC Plant Biol 10, 280, doi:10.1186/1471-2229-10-280 (2010).

32 Yang, H., Lu, P., Wang, Y. & Ma, H. The transcriptome landscape of Arabidopsis male meiocytes from high-throughput sequencing: the complexity and evolution of the meiotic process. Plant J 65, 503–516, doi:10.1111/j.1365-313X.2010.04439.x (2011).

33 Martinez, G., Choudury, S. G. & Slotkin, R. K. tRNA-derived small RNAs target transposable element transcripts. Nucleic Acids Res, doi:10.1093/nar/gkx103 (2017).

34 Wang, X. et al. DNA Methylation Affects Gene Alternative Splicing in Plants: An Example from Rice. Mol Plant 9, 305–307, doi:10.1016/j.molp.2015.09.016 (2016).

35 Regulski, M. et al. The maize methylome influences mRNA splice sites and reveals widespread paramutation-like switches guided by small RNA. Genome Res 23, 1651–1662, doi:10.1101/gr.153510.112 (2013).

36 Naftelberg, S., Schor, I. E., Ast, G. & Kornblihtt, A. R. Regulation of alternative splicing through coupling with transcription and chromatin structure. Annu Rev Biochem 84, 165–198, doi:10.1146/annurev-biochem-060614-034242 (2015).

37 Lev Maor, G., Yearim, A. & Ast, G. The alternative role of DNA methylation in splicing regulation. Trends Genet 31, 274–280, doi:10.1016/j.tig.2015.03.002 (2015).

38 Jiang, H. et al. MULTIPOLAR SPINDLE 1 (MPS1), a novel coiled-coil protein of Arabidopsis thaliana, is required for meiotic spindle organization. Plant J 59, 1001–1010, doi:10.1111/j.1365-313X.2009.03929.x (2009).

39 De Muyt, A. et al. A high throughput genetic screen identifies new early meiotic recombination functions in Arabidopsis thaliana. Plos Genet 5, e1000654, doi:10.1371/journal.pgen.1000654 (2009).

40 Oliver, C., Santos, J. L. & Pradillo, M. Accurate Chromosome Segregation at First Meiotic Division Requires AGO4, a Protein Involved in RNA-Dependent DNA Methylation in Arabidopsis thaliana. Genetics 204, 543–553, doi:10.1534/genetics.116.189217 (2016).

41 Hollister, J. D. & Gaut, B. S. Epigenetic silencing of transposable elements: a trade-off between reduced transposition and deleterious effects on neighboring gene expression. Genome Res 19, 1419–1428, doi:10.1101/gr.091678.109 (2009).

42 Baubec, T., Finke, A., Mittelsten Scheid, O. & Pecinka, A. Meristem-specific expression of epigenetic regulators safeguards transposon silencing in Arabidopsis. EMBO Rep 15, 446–452, doi:10.1002/embr.201337915 (2014).

43 Tran, R. K. et al. DNA methylation profiling identifies CG methylation clusters in Arabidopsis genes. Curr Biol 15, 154–159, doi:10.1016/j.cub.2005.01.008 (2005).

44 Takuno, S. & Gaut, B. S. Body-methylated genes in Arabidopsis thaliana are functionally important and evolve slowly. Mol Biol Evol 29, 219–227, doi:10.1093/molbev/msr188 (2012).

45 Zilberman, D., Gehring, M., Tran, R. K., Ballinger, T. & Henikoff, S. Genome-wide analysis of Arabidopsis thaliana DNA methylation uncovers an interdependence between methylation and transcription. Nat Genet 39, 61–69, doi:10.1038/ng1929 (2007).

## References

46 Coleman-Derr, D. & Zilberman, D. Deposition of histone variant H2A.Z within gene bodies regulates responsive genes. Plos Genet 8, e1002988, doi:10.1371/journal.pgen.1002988 (2012).

47 Stroud, H. et al. Non-CG methylation patterns shape the epigenetic landscape in Arabidopsis. Nat Struct Mol Biol 21, 64–72, doi:10.1038/nsmb.2735 (2014).

48 Zemach, A. et al. The Arabidopsis nucleosome remodeler DDM1 allows DNA methyltransferases to access H1-containing heterochromatin. Cell 153, 193–205, doi:10.1016/j.cell.2013.02.033 (2013).

49 Liang, C., Liu, X., Sun, Y., Yiu, S. M. & Lim, B. L. Global small RNA analysis in fast-growing Arabidopsis thaliana with elevated concentrations of ATP and sugars. BMC Genomics 15, 116, doi:10.1186/1471-2164-15-116 (2014).

50 Martinez, G., Panda, K., Kohler, C. & Slotkin, R. K. Silencing in sperm cells is directed by RNA movement from the surrounding nurse cell. Nat Plants 2, 16030, doi:10.1038/nplants.2016.30 (2016).

51 Armstrong, S. J., Sanchez-Moran, E. & Franklin, F. C. Cytological analysis of Arabidopsis thaliana meiotic chromosomes. Methods Mol Biol 558, 131–145, doi:10.1007/978-1-60761-103-5_9 (2009).

52 Sonobe, S. & Shibaoka, H. Cortical Fine Actin-Filaments in Higher-Plant Cells Visualized by Rhodamine-Phalloidin after Pretreatment with M-Maleimidobenzoyl N-Hydroxysuccinimide Ester. Protoplasma 148, 80–86 (1989).

53 Nakagawa, T. et al. Development of series of gateway binary vectors, pGWBs, for realizing efficient construction of fusion genes for plant transformation. J Biosci Bioeng 104, 34–41, doi:10.1263/jbb.104.34 (2007).

54 Deal, R. B. & Henikoff, S. The INTACT method for cell type-specific gene expression and chromatin profiling in Arabidopsis thaliana. Nat Protoc 6, 56–68, doi:10.1038/nprot.2010.175 (2011).

